# A transcriptomic continuum of differentiation arrest identifies myeloid interface acute leukemias with poor prognosis

**DOI:** 10.1101/2019.12.10.870121

**Authors:** Jonathan Bond, Aleksandra Krzywon, Ludovic Lhermitte, Christophe Roumier, Anne Roggy, Mohamed Belhocine, Alexander Kaden Kheirallah, Patrick Villarese, Guillaume Hypolite, Francine Garnache-Ottou, Sylvie Castaigne, Nicolas Boissel, Vahid Asnafi, Claude Preudhomme, Hervé Dombret, Elisa Laurenti, Elizabeth Macintyre

## Abstract

Classification of acute lymphoblastic and myeloid leukemias (ALL and AML) remains heavily based on phenotypic resemblance to normal hematopoietic precursors. This framework can provide diagnostic challenges for immunophenotypically heterogeneous immature leukemias, and ignores recent advances in understanding of developmental multipotency of diverse normal hematopoietic progenitor populations that are identified by transcriptional signatures. We performed transcriptional analyses of a large series of acute myeloid and lymphoid leukemias and detected significant overlap in gene expression between cases in different diagnostic categories. Bioinformatic classification of leukemias along a continuum of hematopoietic differentiation identified leukemias at the myeloid/T-lymphoid interface, which shared gene expression programs with a series of multi or oligopotent hematopoietic progenitor populations, including the most immature CD34+CD1a-CD7- subset of early thymic precursors. Within these interface acute leukemias (IALs), transcriptional resemblance to early lymphoid progenitor populations and biphenotypic leukemias was more evident in cases originally diagnosed as AML, rather than T-ALL. Further prognostic analyses revealed that expression of IAL transcriptional programs significantly correlated with poor outcome in independent AML patient cohorts. Our results suggest that traditional binary approaches to acute leukemia categorization are reductive, and that identification of IALs could allow better treatment allocation and evaluation of therapeutic options.

## Introduction

Successful management of acute leukemia is underpinned by accurate diagnostic classification, which provides a basis for treatment allocation, risk stratification and implementation of targeted therapies (1). Although knowledge of the molecular landscape of leukemia has increased enormously over the past decades, contemporary classification remains heavily predicated on simple immunophenotypic resemblance to either myeloid or lymphoid normal hematopoietic precursors (2). While this system has historically been successful, some leukemia categories provide specific diagnostic and therapeutic challenges. The current World Health Organization (WHO) classification (2) recognizes acute leukemias of ambiguous lineage that either lack lineage-specific markers (acute undifferentiated leukemias, AUL) or express a mixture of myeloid and lymphoid antigens (mixed phenotype acute leukemias, MPAL). There is little consensus on the best treatment approaches for these patients, and prognosis is usually poor (3–5).

This framework also poses difficulties for some cases of T-acute lymphoblastic leukemia (T-ALL) and acute myeloid leukemia (AML). T-ALL can be subclassified by immunogenotypic and phenotypic resemblance to either immature/ early thymic precursor (ETP), early cortical or late cortical normal T-progenitor equivalents (6, 7). However, the genotypic and phenotypic heterogeneity of immature T-ALLs mean that robust biological classification of this group is not straightforward (8). A subset of these cases harbor mutations that are also commonly seen in AML, suggesting that at least some immature T-ALLs may arise from transformation of a bipotent lympho-myeloid progenitor (9–13). In addition, diagnostic distinction from AML by immunophenotype is often not clear-cut, as immature T-ALLs commonly express myeloid lineage-associated markers (14). Conversely, the most phenotypically immature AML subgroup, M0-AML, is also biologically heterogeneous and expresses lymphoid-associated antigens such as CD7 or TdT in about 50% of cases (15). Immature T-ALLs are frequently chemoresistant and require intensive treatment (10, 14, 16), while M0-AML cases have poor outcomes compared to other AML subgroups (17, 18), so it is clinically important to consider whether improved classification of these cases might allow better therapeutic choices.

Current leukemia classification also takes little account of modern advances in understanding of human hematopoiesis, and the recognition of a diverse range of pluri- and multipotent progenitors, as identified by transcriptional signatures and functional assays (19). In particular, traditional notions of an early lymphoid/myeloid dichotomy have been undermined by the discovery of a multitude of lymphoid committed cell types which retain myeloid potential at different stages of differentiation: within the phenotypic stem cell (20) or progenitor compartment (21–25) and in the thymus (26, 27). The relevance of these cell types in the context of leukemia is only beginning to be explored (22, 28).

Leukemic transcriptome profiling should help to improve categorization, but traditional analytical approaches have their shortcomings. T-ALL can be reproducibly categorized according to a limited number of expression signatures that correlate with the phenotype of differentiation arrest (6, 29, 30). Data may also be interrogated by gene set enrichment analysis (GSEA), which has revealed that immature/ETP-ALLs transcriptionally resemble both normal hematopoietic stem cell (HSC) and immature myeloid precursors (9). However, these approaches rely on comparisons of predefined sample groups, neglect transcriptional heterogeneity of individual leukemias in each group and cannot resolve relationships between groups. These analyses therefore provide limited information about the spectrum of differentiation arrest in acute leukemia.

Evolutions in genomic analytical methods provide an opportunity to refine leukemia classification. We have analyzed a series of acute leukemias that comprised a high proportion of immature T-ALLs and AMLs using several approaches, including the novel Iterative Clustering and Guide Gene Selection method (ICGS). This technique, when applied to single-cell RNA-sequencing data, has been shown to infer cellular states from transcriptional data, identify modules of guide genes that are specific to these cellular developmental states in an unbiased, agnostic manner, and infer developmental relationships between these states (31). We show that application of ICGS to global expression data identifies a continuum of differentiation arrest, which includes a group of myeloid/ T-lymphoid interface leukemias that lack clear lineage identity, and which respond poorly to AML treatment regimens.

## Methods

### Microarray data analysis

All computational analysis was performed in R (v.3.3.2 or above) unless otherwise specified. Data were normalised with *normalize.quantiles* function from the preprocessCore v1.34.0 package and batch effects between 2 independent arrays were corrected using the *ComBat* function (sva package). Hierarchical clustering was performed with the *hclust* function with distance (1-Pearson correlation) and complete clustering method. Principal Component Analysis (PCA) was performed with *prcomp* function. Both hierarchical clustering and PCA were performed on all probes.

### ICGS

ICGS was performed with AltAnalyze software v. 2.1.0 (http://www.altanalyze.org/) using HOPACH clustering, with default settings for gene expression analysis options (moderated t-test for group comparison and Benjamini-Hochberg false discovery rate <0.05). The gene expression filtering option was set to 2. Cell cycle genes were excluded using the most stringent parameter. From the Liu et al. pediatric cohort (32), all samples were used, whereas from the Chen et al. cohort (33) only adult samples (>18 years) were selected. Heatmap visualization of ICGS data was performed in AltAnalyze.

### Differential expression analysis

Differentially expressed genes were derived using the *limma* package (*lmFit* function) for microarray and *DESeq2* for RNA-Seq. Contrast matrices between selected groups are listed in Supplementary Table S1. Genes were considered differentially expressed if Benjamini-Hochberg false discovery rate (FDR) < 0.05. Gene ranking for Gene Set Enrichment Analyis (GSEA) was performed according to *t*-statistic for microarray data or Wald statistic for RNA-seq data. For the thymic subpopulation dataset, most variable genes across all populations were selected as the union of all the probes differentially expressed between any two populations (thymic HVGs, 8751 probes).

### Pathway and Gene Set Enrichment Analysis

GSEA was performed with GSEA software (http://software.broadinstitute.org/gsea/index.jsp) using the C2.all.v6.1 collection of genesets from MSigDB (http://software.broadinstitute.org/gsea/msigdb/index.jsp) or a collection of custom genesets (Supplementary Table S1) derived from datasets generated here or publicly available (19, 23, 24, 34–36). When specific genesets were derived from published data, differential expression analysis was performed as indicated above using the contrasts indicated in Table S1. Differentially expressed genes were then ranked by *t*-statistic for microarray data or by Wald statistic for RNA-seq data and the top 500 genes (or all genes with FDR < 0.05 if <500 genes had FDR<0.05) were selected as genesets to be tested by GSEA. GSEA outputs were either visualised with the EnrichmentMap plugin (FDR Q-value cutoff 0.05) of Cytoscape (v.3.2.0), or with heatmaps generated with Prism software (v.7). ClueGO analysis was performed with the ClueGO plugin (v.2.1.6) of Cytoscape (v.3.2.0), using the GO Term Fusion option and otherwise default parameters.

### Data availability

All gene expression data have been deposited in the GEO portal under the accession numbers GSE131180 (thymic populations isolated from neonatal thymi), GSE131184, GSE131207 (AML and T-ALL samples). All relevant data are also available from the authors.

Other experimental methods are described in the **Supplemental Data**.

## Results

### Transcriptional profiling identifies an AML-like subset of T-ALL

We performed transcriptional profiling of a series of 124 acute T-lymphoid and myeloid leukemias (See Supplementary Methods). The 48 T-ALLs included a high proportion (54.2%) of immature cases, as defined by T-receptor immunogenotype (37), comprising 9 IM0 (germline *TR*), 9 IMD (*TRD* rearrangement only) and 8 IMG (*TRG* and *TRD* rearranged but absent or incomplete *TRB* rearrangement) leukemias. Similarly, 28/76 AML samples (40.8%) were categorized as M0-AML. Patient details are shown in Supplementary Table S2.

Unsupervised hierarchical clustering (HC) analysis of the expression data revealed that T-ALL and AML samples largely formed two distinct groups (HC cluster 1 and HC cluster 2, Figure 1A). Strikingly, 8/48 T-ALLs (16.7%, henceforth ‘AML-like T-ALL’) segregated in the AML cluster in this unsupervised analysis, and clustered together when HC was restricted to T-ALLs (Supplementary Figure S1A). When visualized by Principal Component Analysis (PCA), T-ALL and AML samples were distributed differently along the first principal component. Notably, T-ALL samples clustering with AMLs by HC overlapped with AML samples (Supplementary Figure 1B).

**Figure 1:**
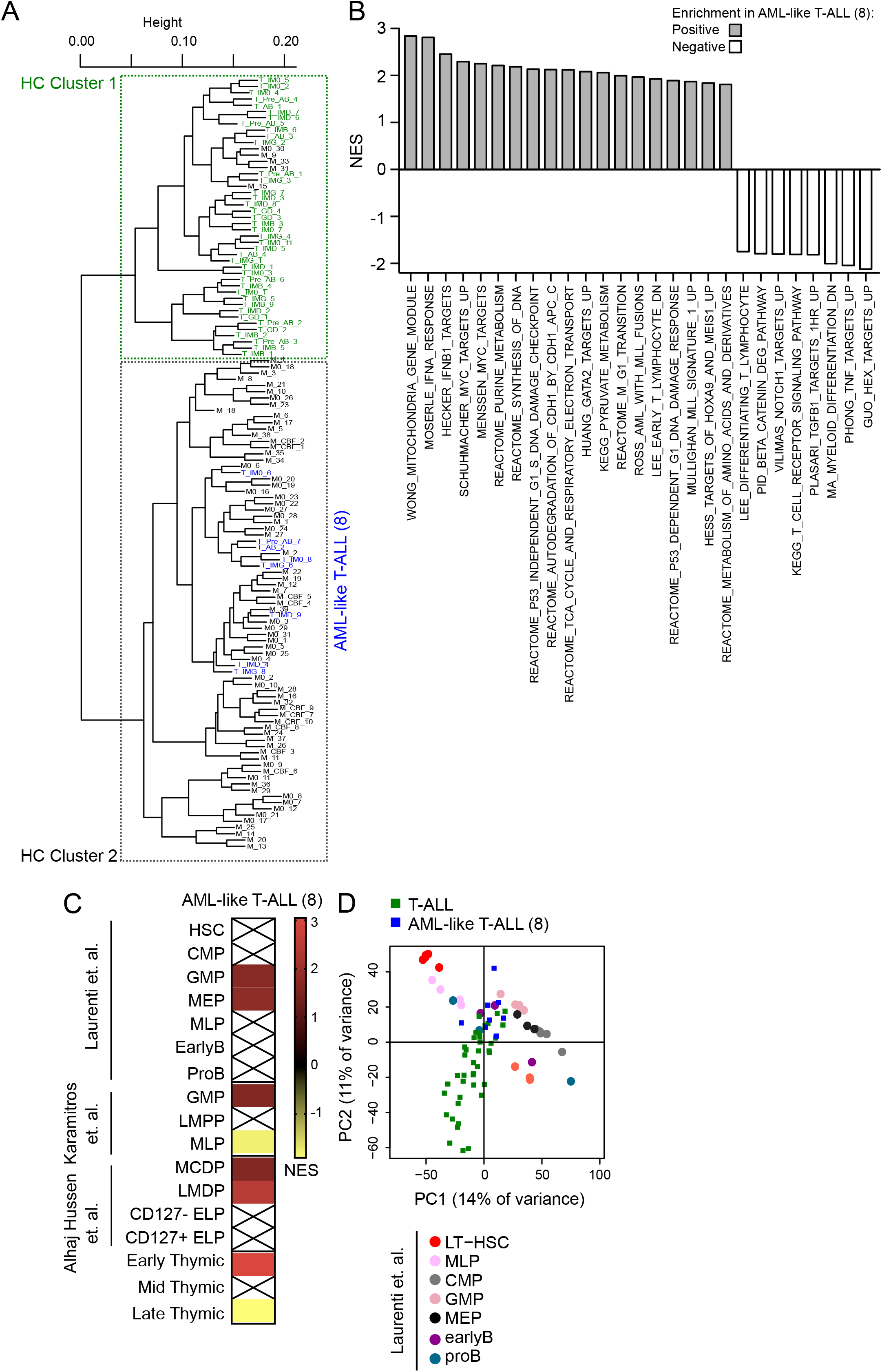
Transcriptional profiling identifies AML-like T-ALLs that are enriched for immature myeloid and thymic progenitor transcriptional signatures. **(A)** Unsupervised hierarchical clustering (HC) of the transcriptional profiles of 124 acute leukemias, comprising 48 T-ALLs and 76 AMLs. A subset of T-ALL cases segregates with the AML cluster. **(B)** GSEA analysis of pathways significantly enriched in AML-like T-ALLs vs the rest of the T-ALL cohort. The MSigDB C2 collection of genesets was used and only selected genesets with FDR < 0.05 are shown. NES = Normalized Enrichment Score. **(C)** Enrichment of selected normal hematopoietic progenitor transcriptional signatures derived from the indicated published datasets or our own analysis of thymic subpopulations (genesets provided in Supplementary Table S4) in AML-like T-ALLs by GSEA. NES = Normalized Enrichment Score, crossed out boxes indicate genesets that are not significantly enriched (FDR > 0.05). HSC = Hematopoietic Stem Cell, CMP = Common Myeloid Progenitor, GMP = Granulocyte-Monocyte Progenitor, MEP = Megakaryocytic-Erythroid Progenitor, MLP = Multi-Lymphoid Progenitor, LMPP = Lymphoid-Primed Multipotent Progenitor, MDCP = Monocyte-Dendritic cell Progenitor, LMDP = Lymphoid-Mono-Dendritic Progenitor, ELP = Early Lymphoid Precursor. **(D)** 2D PCA map of umbilical cord blood stem and progenitor populations and T-ALL gene expression patterns (38); distribution of AML-like T-ALLs (blue squares) is significantly different to that of other T-ALLs (PC1: p= 0.003; PC2: p= 4.1×10^−5^ by two-sided t-test).

Not all of these AML-like T-ALLs exhibited immunogenotypic immaturity (6/8) or had an ETP-ALL immunophenotype (4/7 fully-phenotyped samples) (14), indicating that AML-like transcription features are not restricted to previously identified categories of less differentiated T-ALLs.

### AML-like T-ALL is enriched for myeloid progenitor transcriptional signatures

We next examined the transcriptional differences between AML-like cases and the rest of the T-ALL cohort. 2274 genes (Supplementary Table S3) were significantly differentially expressed between the two groups (FDR <0.05), with 1213 and 1061 respectively upregulated and downregulated in AML-like T-ALLs. Pathway analysis revealed that AML-like T-ALLs had elevated expression of genes involved in cell cycle and mitochondrial, amino-acid and pyruvate metabolism, and high levels of interferon-related genes, MYC, HOXA, MEIS1 and GATA2 targets (Figure 1B). Gene-sets that were previously reported to be upregulated in AML in independent datasets were also significantly over-represented. In contrast, TCR, NOTCH1 and TNF signaling were all downregulated.

We then sought to better characterize AML-like T-ALLs similarity to normal stem and progenitor cells, by performing GSEA using normal umbilical cord blood (UCB) hematopoietic progenitor transcriptional signatures that we previously reported (38). AML-like T-ALLs were significantly enriched for megakaryocytic-erythroid progenitor (MEP) and granulocyte-monocyte progenitor (GMP), but not hematopoietic stem cells (HSC) signatures. These leukemias were also enriched for a GMP signature from an independent data-set (23), and resembled lymphoid-mono-dendritic progenitors (LMDP) from an UCB-derived humanized murine model of early lymphoid development (24) (Figure 1C). To confirm transcriptional similarity to myeloid progenitors, we combined the gene expression of the T-ALL samples with that of highly purified stem and progenitor populations (38) on a 2D PCA map. Consistent with the GSEA results, AML-like T-ALLs localized in the HSPC differentiation space, near GMPs (Figure 1D).

### AML-like T-ALL transcriptionally resembles immature thymic progenitors

While previous analyses of ETP-ALL have evaluated transcriptional proximity to normal ETP cells (9), comprehensive transcriptional comparisons of T-ALL and normal thymic subpopulations are lacking. We performed transcriptional profiling of six phenotypically defined T-lymphoid progenitor groups isolated from a series of human thymi (Supplementary Figure S2A).

The genes most differentially expressed in each subpopulation (Supplementary Figure S2B and Supplementary Table S4) were consistent with known T-lymphopoietic transcriptional patterns. PCA also reflected this developmental progression (Supplementary Figure S2C), which was similar to an *in-vitro* system of human thymocyte differentiation from UCB CD34+ cells (39) (Supplementary Figure S2D).

PCA identified 3 main clusters: a rare (Supplementary Figure S2A) ‘early’ thymic group comprising CD34+CD1a-CD7-samples, a ‘middle’ thymic group comprising CD34+CD1a-CD7+, CD34+CD1a+ and CD4+ ISP samples and a ‘late’ thymic group encompassing the transcriptionally similar CD4+CD8+DP/TRLow and CD4+CD8+DP/TRHigh samples. We derived specific gene expression signatures for each of these clusters and used these in GSEAs to assess the transcriptional similarity of AML-like T-ALLs to normal thymocyte subsets. Strikingly, AML-like T-ALLs were strongly positively enriched for genes that were specifically expressed by the most immature CD34+CD1a-CD7-thymic subpopulation (Figure 1C). Of note, this signature differed from an ETP transcriptional profile that we previously reported, which was derived by comparison to CB stem and progenitor cells (38) (Supplemental Figure 2E-2G). Conversely, when compared with the rest of the T-ALL cohort, AML-like T-ALL samples were negatively enriched for ‘late’ thymic discriminating genes (Figure 1C). Taken together, these results indicate that AML-like T-ALLs share gene expression programs with both UCB-derived myeloid-competent progenitors and the most immature thymic precursors, which also retain myeloid differentiation potential (27).

### Iterative Clustering and Guide Gene Selection analysis identifies a continuum of leukemic differentiation arrest

The recently described ICGS method employs serial iterative clustering with pattern-specific guide genes to define coherent transcriptional patterns between samples and then groups these samples into cellular states that recapitulate developmental trajectories (31). We reasoned this method could help resolve stages of differentiation arrest in leukemia. To test the feasibility of applying this approach to leukemic datasets, we initially used ICGS to analyze two published series of adult (33) and pediatric (32) T-ALL. For both cohorts, the ICGS algorithm unbiasedly identified guide gene modules enriched for human stem and progenitor cells (HSPCs, CD34+), myeloid cells and thymocytes (Supplementary Figure S3A and S3C and Supplementary Table S5), and ordered the T-ALL samples in clusters along a continuum of expression of these genes. Along this spectrum, adult T-ALLs attributed to ICGS clusters with the lowest expression of thymic-associated genes (Groups A and B), but with high expression of HSPC and myeloid genes, were enriched for the ETP-ALL immunophenotype (10, 12–14). For the pediatric cohort (32), ICGS ordering recapitulated in an unsupervised manner the classification the authors had derived linking mutations to thymic developmental stages (Supplementary Figure S3C and S3D). We thus concluded that ICGS allows unbiased classification of leukemic samples according to their stage of differentiation arrest.

We then used ICGS to analyze our patient cohort. ICGS classified these leukemias into five developmental clusters that were defined by the levels of expression of a limited number of guide genes (Figure 2A and Supplementary Table S5) that again predominantly comprised transcripts that discriminate hematopoietic cell types. The proportions of different leukemic phenotypes within each cluster are shown in Figure 2B. Cluster 1 was defined by high expression of thymic- and lymphoid-related genes (e.g. *TCF7*, *LCK*, *BCL11B*), and comprised T-ALL cases exclusively. Conversely, Clusters 4 and 5 were effectively restricted to AML cases, with concentration of Core Binding Factor (CBF)-AMLs in cluster 5. These clusters exhibited increased expression of factors that define myeloid transcriptional modules (e.g. *MPO*, *CEBPE*, *CSF3R*). The intermediate Clusters 2 and 3 were characterized by heterogeneous guide gene expression, and included one third of T-ALL cases (16/48, 33.3%). Notably, the most immature M0 subtype AMLs were predominantly found in these two clusters (24/28, 85.7%), as compared with 14/48 (29.2%, p<0.001 by Fisher test) of non-M0-AML. Also, virtually all AML-like T-ALL samples that were defined by HC (7/8, 87.5%) were found in either Cluster 2 (n=4) or 3 (n=3). ICGS therefore provides a means of classifying leukemias along a spectrum of hematopoietic ontogeny, which in our cohort included a significant number of cases at the interface between T-lymphoid and myeloid lineages. Broadly, these ‘interface’ acute leukemias (IAL) either showed no clear evidence of mature T-lymphoid or mature myeloid identity (Cluster 2), or had a partial HSPC/mature myeloid signature (Cluster 3).

**Figure 2:**
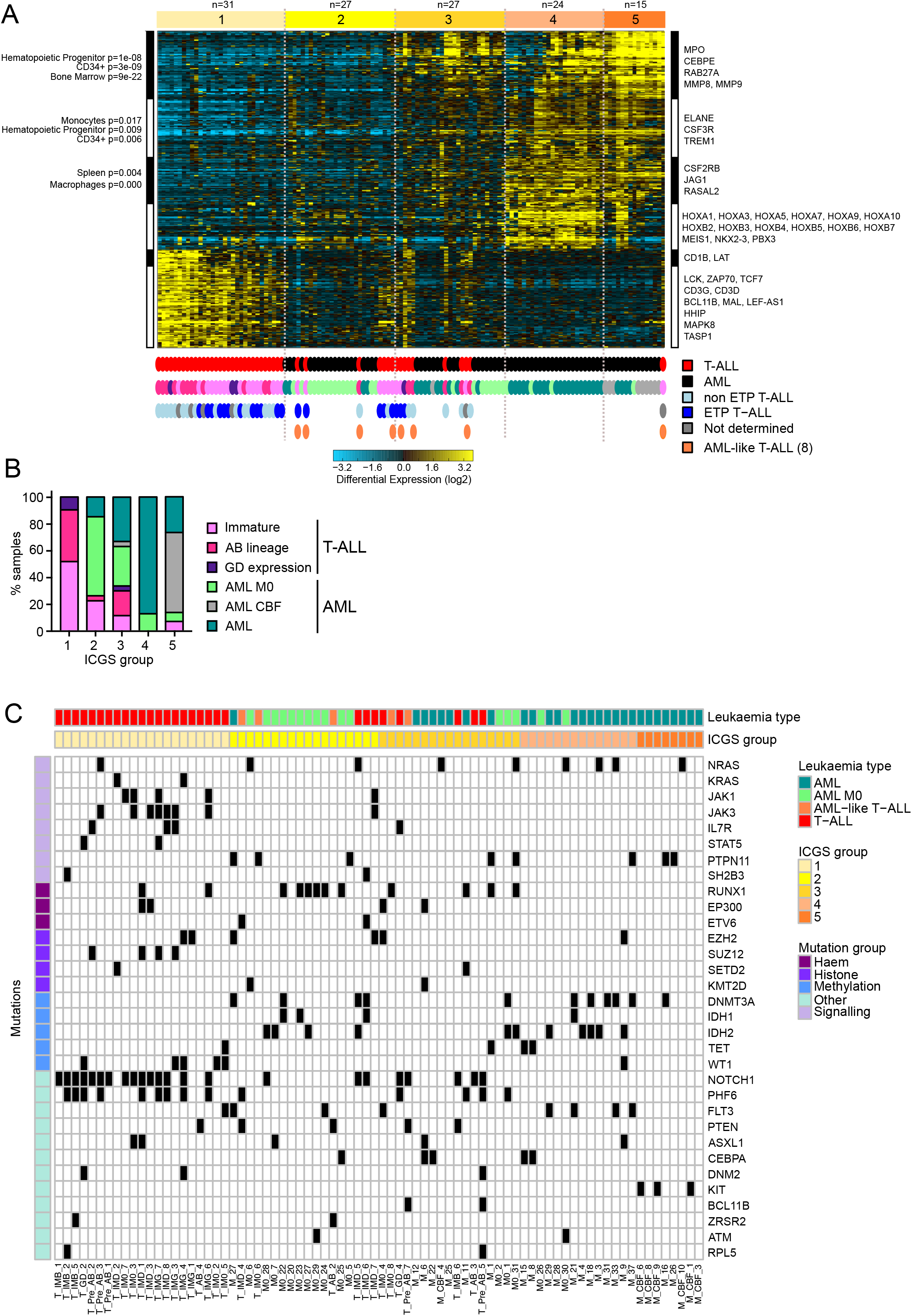
Iterative Clustering and Guide Gene Selection (ICGS) analysis identifies a continuum of leukemic differentiation arrest. **(A-B)** ICGS analysis of adult and pediatric T-ALLs (n=48 samples) and AMLs (n=76 samples) identifies 5 acute leukemia clusters (top). **(A)** Heatmap of expression of guide genes selected by ICGS. Columns represent individual samples. Bars on the top identify ICGS clusters. Rows represent genes, and bars on the side represent blocks of correlated genes. Selected enriched gene ontology groups are shown. Full gene lists are provided in Supplementary Table S5. Leukemic phenotypes are indicated in the bars below the heatmap. **(B)** Proportions of leukemic phenotypic groups in each ICGS cluster. **(C)** Mutations observed in T-ALL (n=34) and AML (n=45) samples ordered according to ICGS analysis in (A). Only mutations found in at least 2 samples are shown.

### Mutational analysis of ICGS-defined clusters

We performed targeted next generation sequencing (NGS) of the 79/124 cases (34 T-ALLs and 45 AMLs) where diagnostic material was available. The NGS panel (Supplementary Table S6) had a predominance of genes that are more often altered in T-ALL, including mutations typically found in the immature subgroup that overlap with those seen in AML (9, 10, 12, 40). Comprehensive results are in Supplementary Table S7, and all mutations detected in ≥ 2 patients are shown in Figure 2C.

Some results were in keeping with the spectrum of differentiation observed. Cluster 1 was enriched for T-ALL type NOTCH pathway-activating mutations (p<0.0001, all comparisons below by Fisher test), while *KIT* mutations correlated with the concentration of CBF-AMLs in Cluster 5 (p=0.0007). However, Cluster 1 was also enriched for mutations in *SUZ12* (p=0.004), *WT1* (p=0.0044) and genes encoding IL7R/JAK/STAT pathway members (p=0.0364), which are normally more frequent in immature T-ALLs (9, 10, 41). Other mutations usually found in less differentiated leukemias (13, 42, 43) were more common in interface cases. Notably, T-ALLs with alterations in DNA methylating factors *DNMT3A*, *IDH1* and *IDH2* (including 4 with double *DNMT3A*/*IDH* mutations) were confined to cluster 2 (p=0.0267). *RUNX1*-mutated AMLs were restricted to interface clusters 2 and 3 (p=0.0015). Surprisingly, AML-like T-ALLs in clusters 1 and 2 had frequent *PTEN* mutations, which are usually found in more differentiated T-ALLs (44). Overall, AML-like T-ALLs were significantly more likely to have *PTEN* mutations than the rest of the T-ALL cases analyzed by NGS (3/6, 50% v 2/28, 7.1%, p=0.0287). Taken together, these results suggest that the spectrum of differentiation arrest defined by ICGS is not directly paralleled by underlying mutational genotype, but may throw light on the stage of arrest associated with well-recognized somatic mutation patterns.

### ICGS identifies myeloid leukemias with early lymphoid transcriptional signatures

Having found that ICGS permits classification of acute leukemias along a spectrum of hematopoietic differentiation, we went on to more precisely characterize the transcriptional identity of individual clusters by GSEA. Analysis of the two published T-ALL cohorts (32, 33) revealed that the least differentiated clusters were enriched for transcriptional signatures from a series of immature myeloid and lymphoid progenitor populations, in addition to HSCs (Supplementary Figure S3F).

Within our cohort, Cluster 1 T-ALLs were strongly enriched for mid- and late-thymic expression profiles, and negatively enriched for both early thymic and UCB HSC and myeloid progenitor signatures. AMLs in Clusters 4 and 5 had broadly converse patterns of positive and negative enrichment (Figure 3A).

**Figure 3:**
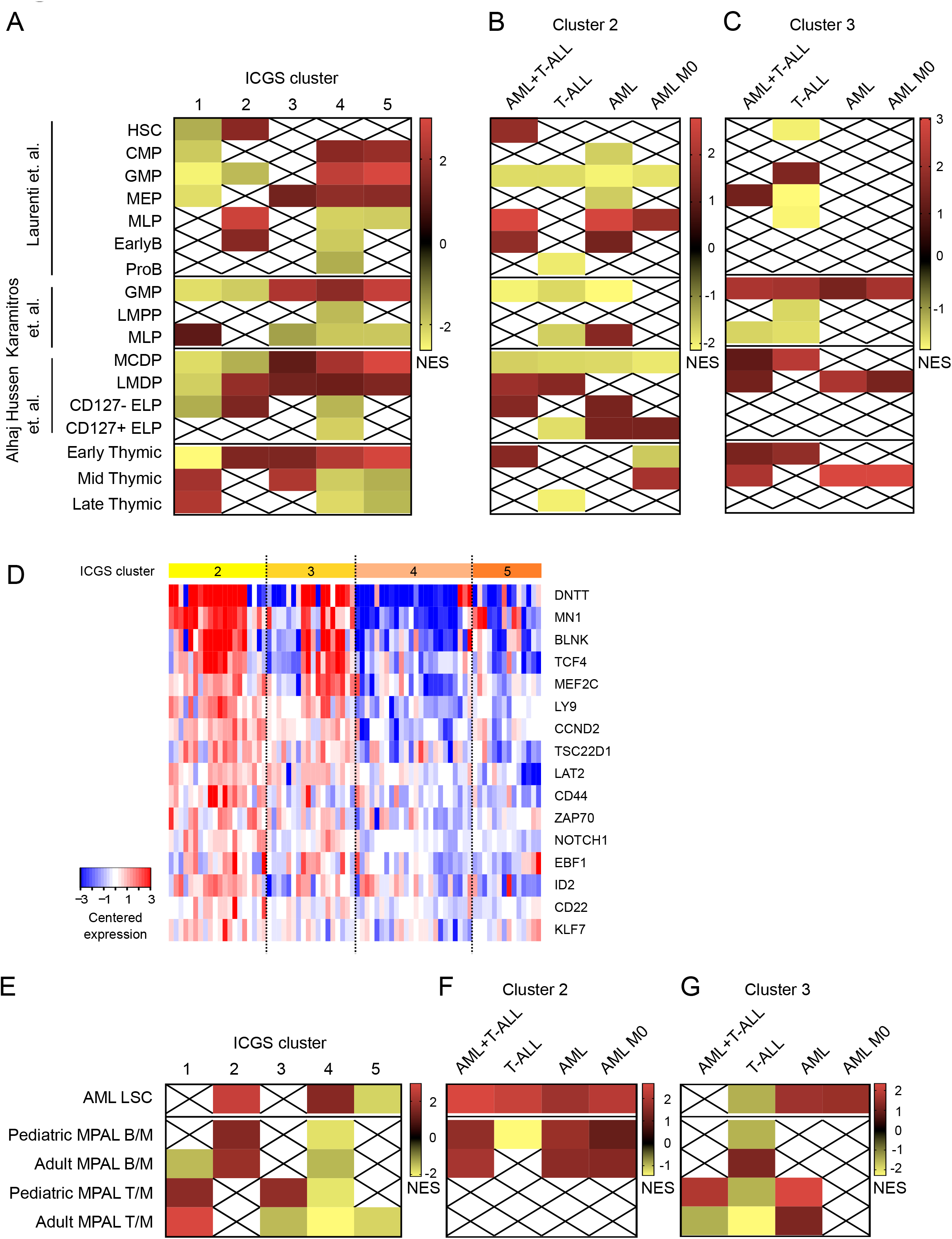
Transcriptional characterization of ICGS-defined clusters. GSEAs using normal hematopoietic precursor transcriptional signatures of **(A)** all clusters, **(B)** interface cluster 2 and **(C)** interface cluster 3. Analyses restricted to either T-ALL, AML-like T-ALL, non-M0-AML and M0-AML are shown. Crossed out boxes indicate genesets that are not significantly enriched (FDR > 0.05). **(D)** Comparison of expression of genes related to B-cell development in interface and non-interface AMLs. **(E) – (G)** Enrichment of leukemic stem cell (LSC) (34) and mixed phenotype acute leukemia (MPAL) (35, 36) transcriptional signatures by GSEA of **(E)** all clusters, **(F)** interface cluster 2 and **(G)** interface cluster 3. Analyses restricted to either T-ALL, AML-like T-ALL, non-M0-AML and M0-AML are shown.

Transcriptional differences in IAL Clusters 2 and 3 were less clear-cut. Cluster 2 IAL (comprising 7 T-ALL, 16 M0-AML and 4 non-M0 AML) were enriched for both HSC and a series of lymphoid progenitor signatures, including MLP, LMDP, early B-cell progenitors, T-oriented CD127-Early Lymphoid Precursors (ELPs) and CD34+CD1a-CD7-early thymic cells (Figure 3A. Cluster 3 cases (9 T-ALL, 8 M0-AML and 10 non-M0-AML) were more likely to be enriched for myeloid profiles (MEP, GMP and UC-derived monocyte-dendritic cell progenitors, MDCP), but also showed transcriptional resemblance to several lymphoid subpopulations, including LMDP and both early and mid-thymic signatures (Figure 3A).

We considered whether this heterogeneity might be driven by differing transcriptional contributions of T-ALLs and AMLs within each cluster. Further analysis of Cluster 2 revealed the surprising finding that while T-ALLs were mostly negatively enriched for lymphoid signatures, AMLs had expression patterns that resembled several lymphoid-competent populations, including MLPs, T-oriented CD127- and B-oriented CD127+ ELPs and early B-cell progenitors (Figure 3B). Similarly, Cluster 3 AMLs showed significant enrichment for LMDP and mid-thymic signatures, while T-ALLs in the same group were more likely to resemble myeloid populations, including GMPs and MDCPs (Figure 3C). These data suggest that interface AMLs demonstrate significant lymphoid orientation, which can be more pronounced than the T-ALLs with which they co-cluster by ICGS. Enrichment for B-lymphoid transcription was particularly evident when expression of genes related to B-cell development was compared in interface and non-interface AMLs (Figure 3D).

### ICGS-defined interface AMLs transcriptionally resemble mixed phenotype leukemia

Further GSEA revealed that interface Cluster 2 was significantly enriched for a myeloid leukemic stem cell (LSC) transcriptional signature (34), and that this enrichment was shared by both T-ALLs and AMLs in this group (Figures 3E and 3F). AMLs in interface Cluster 3 (Figure 3G), and AML-like T-ALLs (NES=1.92; FWER=0.003) were also enriched for the LSC signature, suggesting that expression of leukemia stemness genes is a common feature of IAL cases.

As interface leukemias share expression profiles with a range of progenitors of multipotent lineage capacity, we next tested whether there was any transcriptional similarity to MPALs of either T-lymphoid/myeloid (T/M MPAL) or B-lymphoid/ myeloid (B/M MPAL) phenotype in children (35) and adults (36). We found that interface Clusters 2 and 3 were enriched for B/M MPAL and T/M MPAL signatures respectively, and that enrichment was driven by the AML cases in each group (Figures 3F and 3G). Therefore, in keeping with the results observed in normal progenitor comparisons, transcriptional resemblance to the earliest stages of lymphoid orientation appears to be driven by interface AMLs rather than T-ALLs.

### Interface AMLs have poor outcomes

The fact that interface AMLs exhibit markedly different transcription to other AML cases led us to speculate that these leukemias may have specific biology which in turn might affect clinical behavior. We therefore evaluated the outcome of interface AMLs in two independent studies (45, 46). To identify these cases, we calculated an interface AML (IAL) score based on gene expression differences between interface and non-interface AMLs in our cohort (Supplementary Methods and Table S8). Outcome analyses revealed that AMLs with high IAL scores had significantly shorter survival in both studies (Figures 4A and 4B). Within the ALFA-1701 group, we found that high IAL scores predicted lack of response to gemtuzumab ozogamicin (Figure 4C), which in keeping with our previous results (47), correlated with reduced expression of CD33 in high IAL cases (Figure 4D). Importantly, multivariate analysis of the ALFA-0701 cohort (46) revealed that IAL score predicted outcome independently of other prognostic variables, including cytogenetic classification and the recently described LSC17 score (34) (Table 1). Consistent with this, our IAL signature had almost no overlap with the LSC17 signature, or the extended 48 gene signature that was reported in the same paper (34) (Supplementary Figure S4A and S4B). Full comparison of clinicobiological and mutational profiles of ALFA-0701 patients with high and low IAL scores is shown in Supplementary Table S9. Finally, we evaluated whether IAL High cases had evidence of lymphoid transcriptional activation. In keeping with our earlier results (Figure 3), we found that IAL High cases in both AML cohorts were significantly enriched for both MLP signatures and B-lymphoid gene expression (Supplementary Figure S4C-S4G).

**Figure 4:**
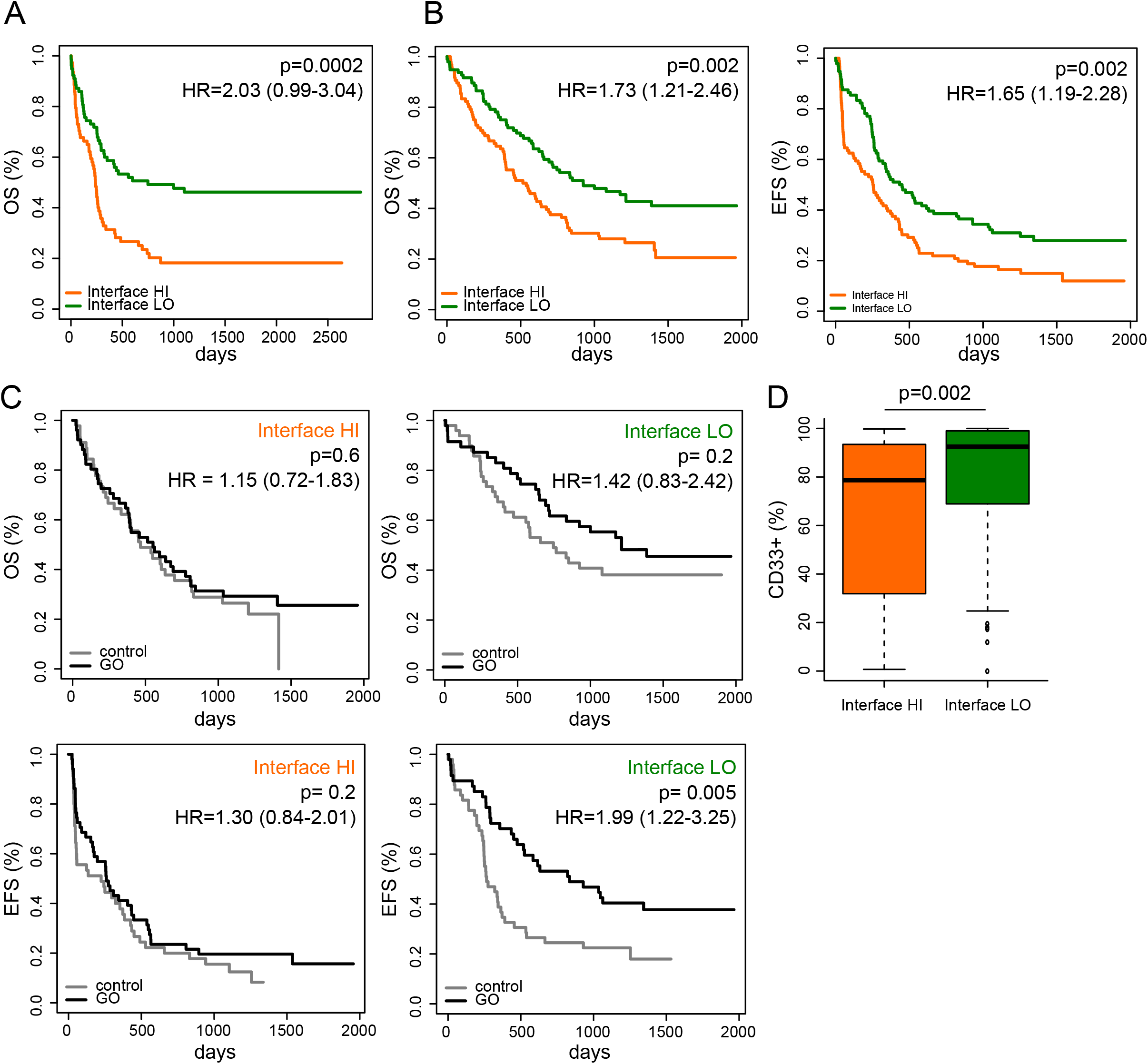
Interface IALs have poor outcomes. Survival comparisons of AMLs with high and low IAL scores in independent cohorts, **(A)** Metzeler *et al* (45) and **(B)** ALFA-1701 (46). OS = Overall Survival. EFS = Event-free Survival. Hazard ratios (HR) and 95% Confidence Intervals for each event and p values are indicated. **(C)** Outcome comparisons according to IAL score and treatment with gemtuzumab ozogamicin (GO) in the ALFA-1701 cohort. **(D)** Comparison of CD33 expression in IAL High and Low cases in the ALFA-1701 cohort. Boxes indicate median, interquartile range and whiskers the 95 percentile.

**Table 1.**
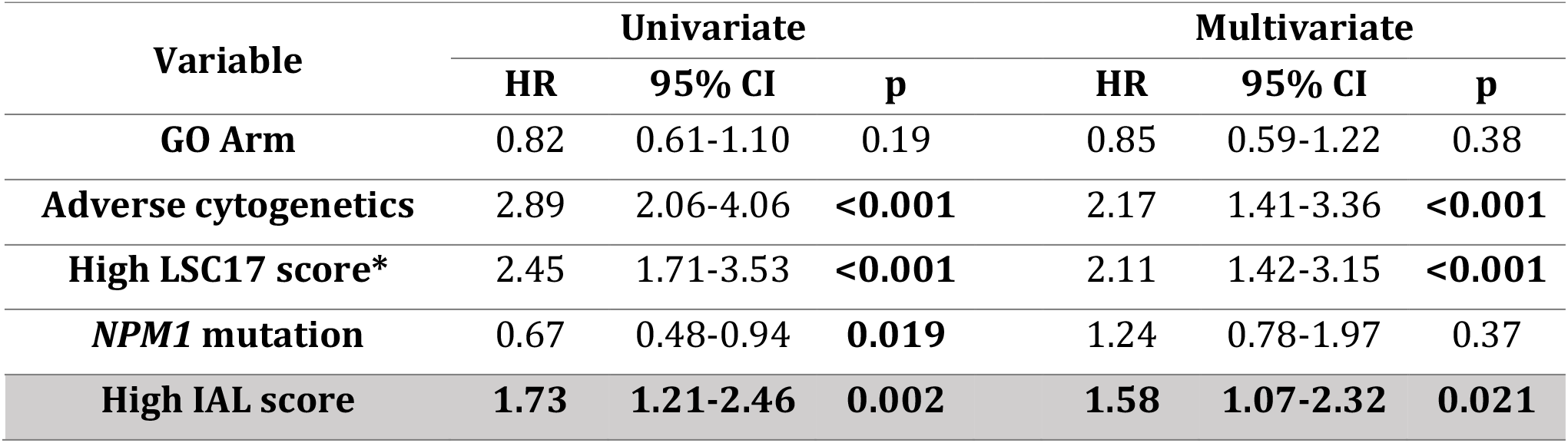
Prognostic impact of IAL score on Overall Survival in the ALFA-0701 trial. Covariates selected for multivariate analyses were selected based on the results of univariate analyses (full results in Supplementary Table S10), with additional retention of GO (gemtuzumab ozogamicin) treatment arm. *The LSC17 score was described in Ng *et al* (34). HR = Hazard Ratio. Statistically significant differences are shown in bold.

## Discussion

In keeping with modern concepts of a hematopoietic progenitor framework that comprises a spectrum of differentiation potential, integrated transcriptional analysis of AMLs and T-ALLs revealed a continuum of leukemic developmental arrest. While AMLs and T-ALLs at either end of the spectrum were specifically enriched for the transcriptional signatures of the corresponding lineage, interface leukemias had evidence of both myeloid and lymphoid precursor gene expression, with early lymphoid signature enrichment being driven by interface AML cases. Specifically, while interface Cluster 3 AMLs had T-lymphoid transcriptional enrichment, interface Cluster 2 AMLs more closely resembled B-oriented lymphoid precursors including early B progenitors, MLPs and CD127+ ELPs (24, 38), and B/Myeloid MPAL (35, 36). This cluster comprised a high proportion of *RUNX1*-mutated M0-AMLs, reported to show B-cell gene activation (48). Overall, these results suggest that these leukemias may be more likely to arise from lymphoid-oriented progenitors and/or be arrested at an early stage of lymphoid orientation (prior to CD19 expression) than is currently recognized.

ICGS clustering presented several important differences with accepted methods of T-ALL categorization by phenotype, immunogenotype or mutational profile (9, 14, 37). For example, the majority of immature T-ALLs defined by *TR* rearrangement (37) (16/26, 61.5%) or ETP-ALL phenotype (12/20, 60%) (14) were in Cluster 1, including those with JAK-STAT pathway mutations (Supplementary Table S7). In addition, IALs had low percentages of *WT1* and *SUZ12* mutations that are typical of ETP-ALLs (9, 10) and positive enrichment for *PTEN* alterations that are more frequent in mature T-ALLs (44, 49). We also noted differences in mutational cooccurrence in these groups. While *PHF6* mutations were always accompanied by *NOTCH1* alterations in Cluster 1, 3/5 *PHF6*-mutated IALs (1/3 T-ALL and 2/2 AML) were *NOTCH1* wild-type. This pattern was also reported in MPAL (35, 50), and suggests that the leukemic phenotype of *PHF6* mutation may correlate with co-expression of other oncogenes, as shown for *TLX3* (51). Interestingly, PHF6 has been shown to regulate B/T lineage plasticity, at least on a *BCR-ABL* leukemic background (52). Interface AMLs were also not restricted to immunophenotypically immature M0 cases, since they included 29% of non-M0 AMLs.

Our description of myeloid/T-lymphoid IALs provides support for recent proposals to define acute myeloid/T-lymphoblastic leukemia (AMTL) as a distinct diagnostic entity (11), but our results also indicate that this group comprises significant molecular and lineage heterogeneity, particularly with regard to lymphoid gene expression. It is also striking that B-lymphoid transcription correlated with poor response to AML treatment regimens. *RUNX1*-mutated AML-M0 cases in our cohort showed B-lymphoid identity, which is consistent with previous reports (48). Intriguingly, *RUNX1-*mutated AMLs have recently been shown to be sensitive to glucocorticoids (53), which form the backbone of ALL induction treatment. Our findings therefore suggest that the poor response of these cases to AML therapy in both adults (54) and children (55) might be improved by better treatment allocation, and would plead against the recent provisional classification of *RUNX1*-mutated AML-M0 with AML (2). Finally, we hope that these data will provide further impetus to include these and other IALs in shared myeloid/lymphoid protocols that might provide better treatment options for patients with these poor-risk leukemias.

## Acknowledgements

The Necker laboratory was supported by the Association Laurette Fugain, La ligue contre le Cancer and the INCa 2007 (DM/FC/sl/RT07) and CARAMELE Translational Research and PhD programs and INCa/AP-HP genetic Plateforme de Ressources Biologiques (PRB). Certain AL samples were collected within the MILE program (56). JB was supported by a Kay Kendall Leukaemia Fund Intermediate Research Fellowship and by the National Children’s Research Centre, Children’s Health Ireland at Crumlin, Dublin. Work in the Laurenti laboratory in Cambridge was supported by a Wellcome/Royal Society Sir Henry Dale Fellowship to EL, the European Hematology Association, BBSRC and by core funding from Wellcome and MRC to the Wellcome-MRC Cambridge Stem Cell Institute. AK was funded by a National Science Centre, Poland research grant (2017/01/X/ST6/01329). We thank Jinyan Huang at the Shanghai Institute of Hematology (SIH) for help in retrieving datasets from the Chinese Leukemia Genotype-Phenotype Archive, as reported by Chen et al (33). We confirm that those who carried out the original analysis and collection of these data at SIH bear no responsibility for our further analysis and interpretation of such data. We thank Koichi Takahashi for annotation of samples from GSE11360. We would also like to thank the Plateforme Biopuces et Séquençage of the IGBMC (Strasbourg) for performance of expression microarray experiments.

## Conflicts of Interest

The authors declare no conflict of interest.

## Notes

#### Summary of Updates

Updated acknowledgments.

https://www.ncbi.nlm.nih.gov/geo/query/acc.cgi?acc=GSE131180

https://www.ncbi.nlm.nih.gov/geo/query/acc.cgi?acc=GSE131207

https://www.ncbi.nlm.nih.gov/geo/query/acc.cgi?acc=GSE131184

