## Supplemental Data for "A transcriptomic continuum of differentiation arrest identifies myeloid interface acute leukemias with poor prognosis"

**Bond *et al***

**Contents:**


- **Supplementary Table Legends**
- **Supplementary Tables S2, S6, S8 - S10**
- **Supplementary Figures S1 - S4**
- **Supplementary Methods**
- **Supplementary References**

**Supplementary Table Legends:**

**Supplementary Table S1:** Gene-sets used for GSEA analysis in this study (see Excel file).

**Supplementary Table S2:** Details of patient cohort.

**Supplementary Table S3:** Differential gene expression analysis comparing AML-like T-ALLs with other T-ALLs (see Excel file). Positive values denote genes with higher expression in AML-like T-ALLs.

**Supplementary Table S4:** Differential gene expression analysis for the comparisons of thymic subset populations indicated in each tab (see Excel files).

**Supplementary Table S5:** ICGS output (see Excel file). First row contains sample names, second row the ICGS clusters, and the following rows guide genes and their normalized expression. The second column indicates the guide genes groups as indicated by the black and white bars in Figure 2A (first tab), Supplementary Figure S3A (second tab (1)) and S3C (third tab (2)).

**Supplementary Table S6:** Genes included in the targeted NGS panel.

**Supplementary Table S7:** Mutational status by NGS (see Excel file). 0 = no mutation, 1 = mutation of known significance, 2 = mutation of unknown significance.

**Supplementary Table S8:** List of genes used for IAL score

**Supplementary Table S9:** Comparison of clinicobiological characteristics and mutational profiles of cases with high and low IAL scores in the ALFA-0701 cohort (3).

**Supplemental Table S10:** Univariate analyses of Overall Survival in the ALFA-0701 cohort (3).

**Supplementary Table S2: Details of the patient cohort.**

Assessment of ETP-ALL phenotype is shown for T-ALL cases only.

| Code | ICGS Cluster | Age | Sex | NGS | ETP phenotype |
|---|---|---|---|---|---|
| T- AB 1 | 1 | 2 | F | No | N/A |
| T- AB 4 | 1 | 18 | F | Yes | No |
| T- GD 1 | 1 | 44 | M | No | N/A |
| T- GD 2 | 1 | 16 | M | Yes | No |
| T- GD 3 | 1 | 45 | M | No | No |
| T- IM0 1 | 1 | 23 | M | No | No |
| T- IM0 11 | 1 | 12 | F | No | Yes |
| T- IM0 2 | 1 | 8 | F | Yes | Yes |
| T- IM0 3 | 1 | 40 | M | Yes | Yes |
| T- IM0 5 | 1 | 22 | M | Yes | Yes |
| T- IM0 7 | 1 | 43 | M | Yes | Yes |
| T- IMB 1 | 1 | 5 | M | Yes | No |
| T- IMB 2 | 1 | 28 | M | Yes | No |
| T- IMB 3 | 1 | 22 | M | No | No |
| T- IMB 4 | 1 | 5 | M | No | No |
| T- IMB 5 | 1 | 52 | M | Yes | No |
| T- IMB 9 | 1 | 10 | M | No | N/A |
| T- IMD 1 | 1 | 37 | M | Yes | No |
| T- IMD 2 | 1 | 28 | M | Yes | Yes |
| T- IMD 3 | 1 | 29 | M | Yes | Yes |
| T- IMD 8 | 1 | 41 | M | Yes | Yes |
| T- IMG 1 | 1 | 23 | M | Yes | Yes |
| T- IMG 3 | 1 | 40 | F | Yes | Yes |
| T- IMG 4 | 1 | 35 | M | Yes | Yes |
| T- IMG 5 | 1 | 15 | F | No | N/A |
| T- IMG 6 | 1 | 31 | M | Yes | No |
| T- IMG 7 | 1 | 33 | M | Yes | Yes |
| T- Pre-ab 1 | 1 | 20 | M | Yes | No |
| T- Pre-ab 2 | 1 | 10 | M | Yes | No |
| T- Pre-ab 3 | 1 | 34 | F | Yes | No |
| T- Pre-ab 6 | 1 | 28 | M | No | No |
| M- 17 | 2 | 26 | F | No | |
| M- 2 | 2 | 7 | M | No | |
| M- 27 | 2 | 60 | F | Yes | |
| M- 5 | 2 | 59 | F | No | |
| M0- 16 | 2 | 36 | F | No | |
| M0- 17 | 2 | 68 | M | No | |
| M0- 19 | 2 | 56 | F | No | |
| M0- 20 | 2 | 49 | M | Yes | |
| M0- 21 | 2 | 44 | F | No | |
| M0- 22 | 2 | 74 | M | Yes | |
| M0- 23 | 2 | 79 | M | Yes | |
| M0- 24 | 2 | 63 | M | Yes | |
| M0- 25 | 2 | 62 | M | Yes | |
| M0- 27 | 2 | 76 | M | Yes | |
| M0- 28 | 2 | 19 | M | Yes | |
| M0- 29 | 2 | 39 | F | Yes | |
| M0- 5 | 2 | 31 | M | Yes | |
| M0- 6 | 2 | 77 | F | Yes | |
| M0- 7 | 2 | | M | Yes | |
| M0- 9 | 2 | | M | No | |
| T- AB 2 | 2 | 17 | M | Yes | No |
| T- IM0 6 | 2 | 10 | M | Yes | Yes |
| T- IMD 4 | 2 | 28 | M | Yes | Yes |
| T- IMD 5 | 2 | 31 | M | Yes | Yes |
| T- IMD 6 | 2 | 56 | M | Yes | Yes |
| T- IMD 7 | 2 | 27 | M | Yes | No |
| T- IMG 8 | 2 | 25 | M | No | Yes |
| M- 1 | 3 | 57 | F | Yes | |
| M- 11 | 3 | 46 | F | Yes | |
| M- 12 | 3 | 13 | M | Yes | |
| M- 19 | 3 | 47 | M | No | |

| Code | ICGS Cluster | Age | Sex | NGS | ETP phenotype |
|---|---|---|---|---|---|
| M- 22 | 3 | 56 | F | Yes | |
| M- 25 | 3 | 17 | F | Yes | |
| M- 39 | 3 | 51 | M | No | |
| M- 6 | 3 | 49 | M | Yes | |
| M- 7 | 3 | 59 | M | No | |
| M- CBF 4 | 3 | 49 | M | Yes | |
| M0- 1 | 3 | 62 | F | Yes | |
| M0- 10 | 3 | 23 | F | No | |
| M0- 12 | 3 | 68 | M | No | |
| M0- 2 | 3 | 27 | F | Yes | |
| M0- 3 | 3 | 17 | M | Yes | |
| M0- 31 | 3 | 56 | M | Yes | |
| M0- 4 | 3 | 9 | F | No | |
| M0- 8 | 3 | | M | No | |
| T- AB 3 | 3 | 22 | M | Yes | No |
| T- GD 4 | 3 | 46 | M | Yes | Yes |
| T- IM0 4 | 3 | 30 | M | Yes | Yes |
| T- IM0 8 | 3 | 7 | M | Yes | Yes |
| T- IMB 6 | 3 | 20 | M | Yes | No |
| T- IMD 9 | 3 | 7 | F | No | N/A |
| T- Pre-ab 4 | 3 | 26 | F | No | No |
| T- Pre-ab 5 | 3 | 42 | M | Yes | No |
| T- Pre-ab 7 | 3 | 25 | M | Yes | No |
| M- 10 | 4 | 44 | F | No | |
| M- 13 | 4 | 48 | F | No | |
| M- 15 | 4 | 51 | M | Yes | |
| M- 18 | 4 | 59 | F | Yes | |
| M- 21 | 4 | 43 | F | Yes | |
| M- 23 | 4 | 29 | F | No | |
| M- 24 | 4 | 27 | M | No | |
| M- 28 | 4 | 48 | M | Yes | |
| M- 29 | 4 | 38 | M | Yes | |
| M- 3 | 4 | 61 | M | Yes | |
| M- 31 | 4 | 50 | F | Yes | |
| M- 32 | 4 | 17 | M | No | |
| M- 33 | 4 | 46 | F | Yes | |
| M- 34 | 4 | 25 | F | No | |
| M- 35 | 4 | 47 | F | No | |
| M- 36 | 4 | 39 | F | No | |
| M- 37 | 4 | 30 | M | Yes | |
| M- 38 | 4 | 39 | M | No | |
| M- 4 | 4 | 43 | M | Yes | |
| M- 8 | 4 | 50 | F | Yes | |
| M- 9 | 4 | 41 | F | Yes | |
| M0- 18 | 4 | 49 | F | No | |
| M0- 26 | 4 | 56 | F | Yes | |
| M0- 30 | 4 | 18 | M | Yes | |
| M- 14 | 5 | 41 | F | No | |
| M- 16 | 5 | 58 | F | Yes | |
| M- 20 | 5 | 35 | M | No | |
| M- 26 | 5 | 25 | M | Yes | |
| M- CBF 1 | 5 | 59 | F | Yes | |
| M- CBF 10 | 5 | 14 | M | Yes | |
| M- CBF 2 | 5 | 62 | F | No | |
| M- CBF 3 | 5 | 67 | M | Yes | |
| M- CBF 5 | 5 | 14 | F | No | |
| M- CBF 6 | 5 | 63 | M | Yes | |
| M- CBF 7 | 5 | 49 | F | No | |
| M- CBF 8 | 5 | 29 | M | Yes | |
| M- CBF 9 | 5 | 54 | M | Yes | |
| M0- 11 | 5 | 83 | M | No | |
| T- IMG 2 | 5 | 5 | F | No | |

### Supplementary Table S6: Genes included in the targeted NGS panel.

| Gene | Transcript | CCDS | Description |
|---|---|---|---|
| *AKT1* | ENST00000554581 | CCDS9994 | v-akt murine thymoma viral oncogene homolog 1 |
| *ASXL1* | ENST00000375687 | CCDS13201 | additional sex combs like 1 (Drosophila) |
| *ATM* | ENST00000278616 | CCDS31669 | ataxia telangiectasia mutated |
| *BCL11B* | ENST00000345514 | CCDS9949 | B-cell CLL/lymphoma 11B (zinc finger protein) |
| *BCOR* | ENST00000342274 | CCDS14250 | BCL6 corepressor |
| *CARD11* | ENST00000396946 | CCDS5336 | caspase recruitment domain family, member 11 |
| *CCR4* | ENST00000330953 | CCDS2656 | chemokine (C-C motif) receptor 4 |
| *CD58* | ENST00000457047 | CCDS44199 | CD58 molecule |
| *CEBPA* | ENST00000498907 | CCDS54243 | CCAAT/enhancer binding protein (C/EBP), alpha |
| *CNOT3* | ENST00000406403 | CCDS12880 | CCR4-NOT transcription complex, subunit 3 |
| *CSNK1A1* | ENST00000377843 | CCDS47303 | casein kinase 1, alpha 1 |
| *CTCF* | ENST00000264010 | CCDS10841 | CCCTC-binding factor (zinc finger protein) |
| *CUL3* | ENST00000264414 | CCDS2462 | cullin 3 |
| *CXCR4* | ENST00000409817 | CCDS33295 | chemokine (C-X-C motif) receptor 4 |
| *DDX3X* | ENST00000399959 | CCDS43931 | DEAD (Asp-Glu-Ala-Asp) box helicase 3, X-linked |
| *DNM2* | ENST00000359692 | CCDS32907 | dynamin 2 |
| *DNMT3A* | ENST00000264709 | CCDS33157 | DNA (cytosine-5-)-methyltransferase 3 alpha |
| *EED* | ENST00000263360 | CCDS8273 | embryonic ectoderm development |
| *EP300* | ENST00000263253 | CCDS14010 | E1A binding protein p300 |
| *ETV6* | ENST00000396373 | CCDS8643 | ets variant 6 |
| *EZH2* | ENST00000320356 | CCDS5891 | enhancer of zeste homolog 2 (Drosophila) |
| *FAS* | ENST00000355740 | CCDS7393 | Fas cell surface death receptor |
| *FBXW7* | ENST00000263981 | CCDS3778 | F-box and WD repeat domain containing 7, E3 ubiquitin protein ligase |
| *FLT3* | ENST00000241453 | CCDS31953 | fms-related tyrosine kinase 3 |
| *FYN* | ENST00000354650 | CCDS5094 | FYN oncogene related to SRC, FGR, YES |
| *GATA3* | ENST00000379328 | CCDS31143 | GATA binding protein 3 |
| *HACE1* | ENST00000262903 | CCDS5050 | HECT domain and ankyrin repeat containing E3 ubiquitin protein ligase 1 |
| *HNRNPA2B1* | ENST00000356674 | CCDS5397 | heterogeneous nuclear ribonucleoprotein A2/B1 |
| *HRAS* | ENST00000417302 | CCDS7699 | Harvey rat sarcoma viral oncogene homolog |
| *IDH1* | ENST00000415913 | CCDS2381 | isocitrate dehydrogenase 1 (NADP+), soluble |
| *IDH2* | ENST00000330062 | CCDS10359 | isocitrate dehydrogenase 2 (NADP+), mitochondrial |
| *IKZF1* | ENST00000349824 | CCDS69299 | IKAROS family zinc finger 1 (Ikaros) |
| *IL7R* | ENST00000303115 | CCDS3911 | interleukin 7 receptor |
| *IRF4* | ENST00000380956 | CCDS4469 | interferon regulatory factor 4 |
| *JAK1* | ENST00000342505 | CCDS41346 | Janus kinase 1 |
| *JAK3* | ENST00000458235 | CCDS12366 | Janus kinase 3 |
| *KIT* | ENST00000288135 | CCDS3496 | v-kit Hardy-Zuckerman 4 feline sarcoma viral oncogene homolog |
| *KMT2A* | ENST00000534358 | CCDS55791 | lysine (K)-specific methyltransferase 2A |
| *KMT2D* | ENST00000301067 | CCDS44873 | lysine (K)-specific methyltransferase 2D |
| *KRAS* | ENST00000311936 | CCDS8702 | Kirsten rat sarcoma viral oncogene homolog |
| *LEF1* | ENST00000379951 | CCDS47122 | lymphoid enhancer-binding factor 1 |
| *NF1* | ENST00000358273 | CCDS42292 | neurofibromin 1 |
| *NOTCH1* | ENST00000277541 | CCDS43905 | notch 1 |
| *NRAS* | ENST00000369535 | CCDS877 | neuroblastoma RAS viral (v-ras) oncogene homolog |
| *PHF6* | ENST00000332070 | CCDS14639 | PHD finger protein 6 |
| *PIK3CA* | ENST00000263967 | CCDS43171 | phosphatidylinositol-4,5-bisphosphate 3-kinase, catalytic subunit alpha |
| *PIK3R1* | ENST00000521381 | CCDS3993 | phosphoinositide-3-kinase, regulatory subunit 1 (alpha) |
| *POT1* | ENST00000357628 | CCDS5793 | protection of telomeres 1 |
| *PTEN* | ENST00000371953 | CCDS31238 | phosphatase and tensin homolog |
| *PTPN11* | ENST00000351677 | CCDS9163 | protein tyrosine phosphatase, non-receptor type 11 |
| *PTPN6* | ENST00000456013 | CCDS44821 | protein tyrosine phosphatase, non-receptor type 6 |
| *PTPRD* | ENST00000381196 | CCDS43786 | protein tyrosine phosphatase, receptor type, D |
| *RB1* | ENST00000267163 | CCDS31973 | retinoblastoma 1 |
| *RELN* | ENST00000428762 | CCDS47680 | reelin |
| *RHOA* | ENST00000418115 | CCDS2795 | ras homolog family member A |
| *RPL10* | ENST00000424325 | CCDS14746 | ribosomal protein L10 |
| *RPL5* | ENST00000370321 | CCDS741 | ribosomal protein L5 |
| *RUNX1* | ENST00000344691 | CCDS42922 | runt-related transcription factor 1 |
| *SETD2* | ENST00000409792 | CCDS2749 | SET domain containing 2 |
| *SF3B1* | ENST00000335508 | CCDS33356 | splicing factor 3b, subunit 1, 155kDa |
| *SH2B3* | ENST00000341259 | CCDS9153 | SH2B adaptor protein 3 |
| *STAT3* | ENST00000264657 | CCDS32656 | signal transducer and activator of transcription 3 (acute-phase response factor) |
| *STAT5B* | ENST00000293328 | CCDS11423 | signal transducer and activator of transcription 5B |
| *SUZ12* | ENST00000322652 | CCDS11270 | SUZ12 polycomb repressive complex 2 subunit |
| *TAL1* | ENST00000294339 | CCDS547 | T-cell acute lymphocytic leukemia 1 |
| *TBL1XR1* | ENST00000457928 | CCDS46961 | transducin (beta)-like 1 X-linked receptor 1 |
| *TDRD6* | ENST00000544460 | CCDS55017 | tudor domain containing 6 |
| *TET2* | ENST00000540549 | CCDS47120 | tet methylcytosine dioxygenase 2 |
| *TET3* | ENST00000409262 | CCDS46339 | tet methylcytosine dioxygenase 3 |
| *TP53* | ENST00000420246 | CCDS45606 | tumor protein p53 |
| *WT1* | ENST00000332351 | CCDS7878 | Wilms tumor 1 |
| *ZEB1* | ENST00000446923 | CCDS44370 | zinc finger E-box binding homeobox 1 |
| *ZRSR2* | ENST00000307771 | CCDS14172 | zinc finger (CCCH type), RNA-binding motif and serine/arginine rich 2 |

**Supplementary Table S8: List of genes used for IAL score**.

| | |
|---|---|
| *CD34* | *MAN1A1* |
| *LDLRAD4* | *LY9* |
| *ATP10A* | *RBM8A* |
| *TSPAN7* | *DYRK3* |
| *STARD9* | *GUCY1A3* |
| *SMAD1* | *KAT6A* |
| *BAALC* | *TAB2* |
| *ZMIZ1* | *MBTD1* |
| *LOC105373495* | *STT3B* |
| *KMT2A* | *CD109* |
| *NPR3* | *PRKACB* |
| *MLLT3* | *CELF2* |
| *BBX* | *UBXN4* |
| *SLC9A7* | *PRKCE* |
| *FOXN3* | *CEP68* |
| *MN1* | *SLC25A30* |
| *PSIP1* | *SHANK3* |
| *SDK2* | *SETBP1* |
| *AFF3* | *AKAP2* |
| *RPL13A* | *ITGA4* |
| *UPF2* | *SMURF2* |
| *GBP4* | *ATP2B4* |
| *ITM2C* | *ZNF251* |
| *FLJ10038* | *MTHFD1* |
| *IFI16* | *SPRY1* |
| *C5orf56* | *RPS6* |
| *MEF2A* | *ELK4* |
| *AKT3* | *PRR5L* |
| *KIAA0754* | *C2CD2* |
| *CD200* | *TLK1* |
| *CLSTN1* | *RNF125* |
| *B4GALT6* | *DDHD1* |
| *KIAA0125* | *UBASH3B* |
| *CD2AP* | *CHRDL1* |
| *TMTC4* | *NCOA7* |
| *GNB5* | *LOC646778* |
| *KLHL13* | *GLS* |
| *TTC3* | *SUMO4* |
| *ABHD17B* | *SLC38A1* |
| *DCK* | *GNG7* |
| *PROM1* | *HMGB1* |
| *PIK3C2B* | *TSPAN5* |
| *ABCB1* | *SH3RF1* |
| *LINC01181* | *PTAR1* |
| *XYLT1* | *GPATCH11* |
| *SPTBN1* | *SEC61A2* |
| *SLC38A2* | *PROSER2* |
| *KATNAL1* | *NOL4L* |
| *GNAI1* | *INPP4B* |
| *RUNX3* | *ERG* |

**Supplementary Table S9: Comparison of clinicobiological characteristics and mutational profiles of cases with high and low IAL scores in the ALFA-0701 cohort (3).**

| | Low IAL score | High IAL score | P values |
|---|---|---|---|
| **Patients, N** | 96 | 96 | - |
| **Median age, years (range)** | 62.0 years (50-70) | 62.4 years (50-70) | 0.55 |
| **WBC, G/L (range)** | 6.9 (0.15-187) | 4.9 (0.5-211) | 0.68 |
| **CD33 expression <70%, N/tested (%)** | 17/67 | 31/70 | 0.031 |
| **Cytogenetic risk, N (%)** | | | 0.003 |
| Favorable | 1 (1%) | 3 (3%) | - |
| Intermediate | 72 (75%) | 57 (59.5%) | - |
| Adverse | 12 (12.5%) | 31 (32.5%) | 0.002 |
| NA | 11 (11.5%) | 5 (5%) | - |
| **ELN RISK, N (%)** | - | - | 0.005 |
| Favorable | 22 | 14 | - |
| Intermediate | 51 | 46 | - |
| Adverse | 12 | 31 | 0.002 |
| NA | 11 | 5 | - |
| **High LSC17 score, N (%)** | 36 (37.5%) | 60 (62.5%) | <0.001 |
| **GENE MUTATIONS, N mutated/tested (%)** | - | - | - |
| *NPM1* | 43/96 | 17/94 | <0.001 |
| *FLT3-ITD* | 19/96 | 14/95 | 0.45 |
| *IDH1* | 8/85 | 12/84 | 0.35 |
| *IDH2* | 13/85 | 11/84 | 0.83 |
| *DNMT3A* | 26/85 | 20/84 | 0.39 |
| *TET2* | 15/85 | 12/84 | 0.68 |
| *WT1* | 3/85 | 6/84 | 0.33 |
| *ASXL1* | 8/85 | 10/84 | 0.63 |
| *RUNX1* | 6/85 | 17/84 | 0.014 |
| sAML-type gene mutations * | 20/85 | 38/84 | 0.004 |

*sAML = secondary AML type mutations, including *ASXL1, SRSF2, STAG2, BCOR, U2AF1, EZH2, SF3B1* and/or *ZRSR2* (Lindsley *et al*. Blood 2015) (4)

**Supplemental Table S10: Univariate analyses of Overall Survival in the ALFA-0701 cohort (3).**

| Variable | Patients, N | HR | 95% CI | P values |
|---|---|---|---|---|
| **GO arm** | 278 | 0.82 | 0.61-1.10 | 0.19 |
| **Age (continuous variable)** | 278 | 1.02 | 0.99-1.05 | 0.16 |
| **WBC (continuous variable)** | 277 | 1.003 | 0.99-1.01 | 0.071 |
| **Adverse cytogenetics** | 249* | 2.89 | 2.06-4.06 | **<0.001** |
| **High CD33 expression (≥70%)** | 200 | 0.86 | 0.60-1.23 | 0.41 |
| **High LSC17 score** | 192 | 2.45 | 1.71-3.53 | **<0.001** |
| ***NPM1* mutation** | 274 | 0.67 | 0.48-0.94 | 0.019 |
| ***FLT3*-ITD mutation** | 275 | 1.06 | 0.72-1.57 | 0.76 |
| ***RUNX1* mutation** | 232 | 1.11 | 0.73-1.70 | 0.62 |
| **sAML-type gene mutations#** | 232 | 1.17 | 0.85-1.62 | 0.34 |
| **High IAL score** | **192** | **1.73** | **1.21-2.46** | **0.002** |

GO = Gemtuzumab Ozogamicin.

*Patients with cytogenetic failure were excluded, leaving 58 adverse and 191 favorable/intermediate cases for analysis.

#sAML = secondary AML type mutations, including *ASXL1, SRSF2, STAG2, BCOR, U2AF1, EZH2, SF3B1* and/or *ZRSR2* (Lindsley *et al*. Blood 2015) (4).

Covariates with significant differences (highlighted in bold) were selected for multivariate analyses, with additional retention of GO treatment arm (Table 1 in main manuscript).

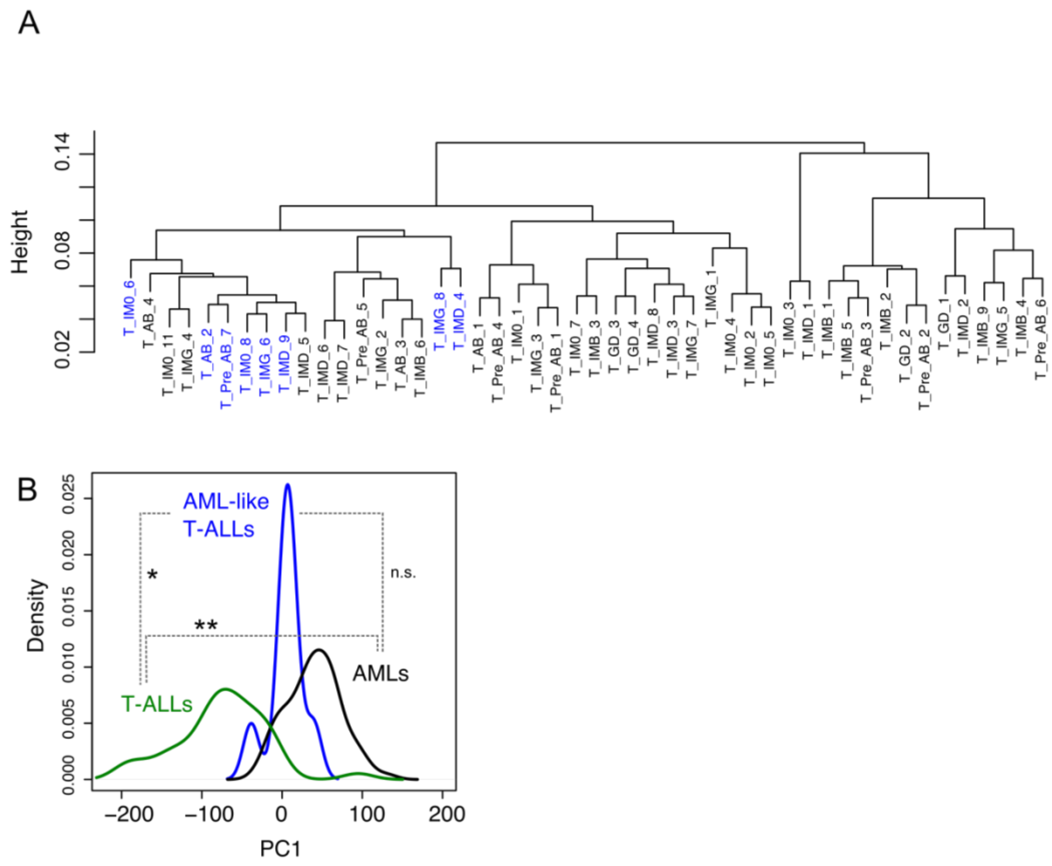


**Supplementary Figure S1: Hierarchical clustering and Principal Component Analysis. (A)** Unsupervised hierarchical clustering (HC) of the transcriptional profiles of the 48 T-ALL samples in the patient cohort. AML-like cases identified by HC in Figure 1A are indicated in blue. **(B)** Principal Component Analysis (PCA) of the sample cohort of T-ALLs and AMLs. Density of distribution of samples in each group along PC1 is indicated. *: $p<0.05$ and **: $p<0.01$ by Kruskal-Wallis test.

A

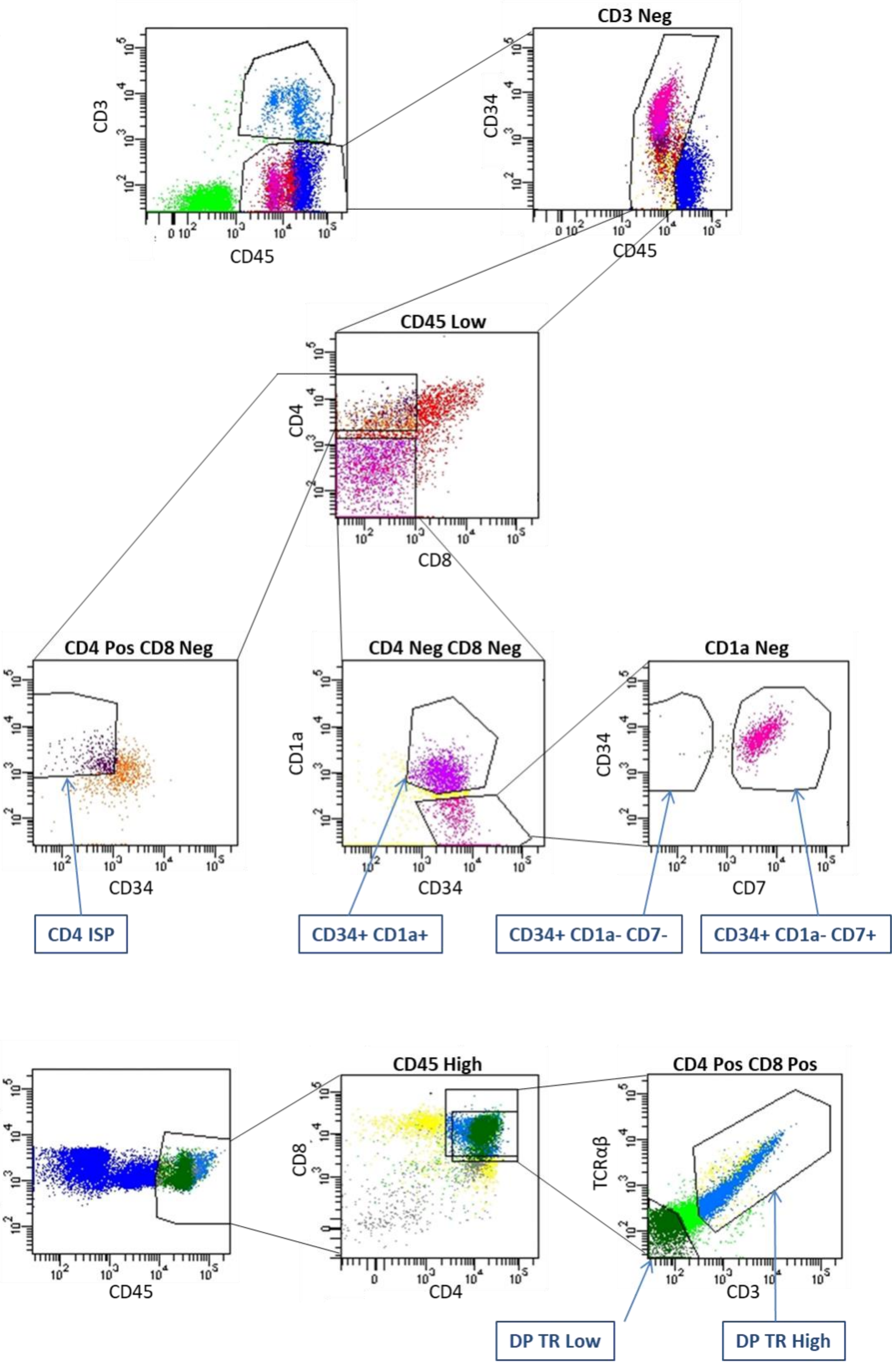

CD3
CD45
CD3 Neg
CD34
CD45
CD45 Low
CD4
CD8
CD4 Pos CD8 Neg
CD1a
CD34
CD4 Neg CD8 Neg
CD1a
CD34
CD1a Neg
CD34
CD7
CD4 ISP
CD34+ CD1a+
CD34+ CD1a- CD7-
CD34+ CD1a- CD7+
SSC
CD45
CD45 High
CD8
CD4
CD4 Pos CD8 Pos
TCRαβ
CD3
DP TR Low
DP TR High

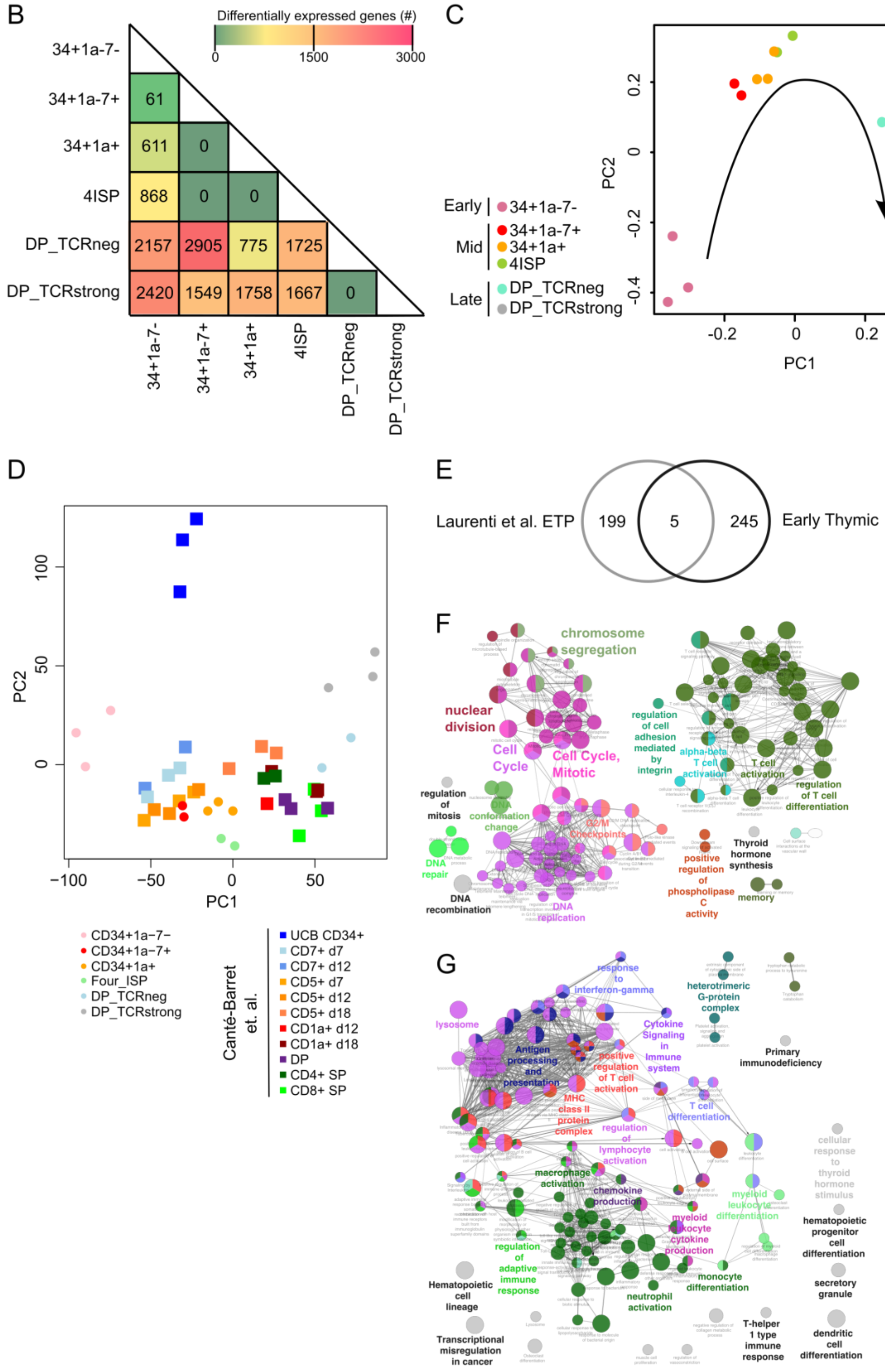
B
Differentially expressed genes (#)
0
1500
3000
34+1a-7-
34+1a-7+
34+1a+
4ISP
DP_TCRneg
DP_TCRstrong
61
611
0
868
0
0
2157
2905
775
1725
2420
1549
1758
1667
0
C
PC2
PC1
Early
Mid
Late
34+1a-7-
34+1a-7+
34+1a+
4ISP
DP_TCRneg
DP_TCRstrong
D
PC2
PC1
CD34+1a−7−
CD34+1a−7+
CD34+1a+
Four_ISP
DP_TCRneg
DP_TCRstrong
Canté-Barret et. al.
UCB CD34+
CD7+ d7
CD7+ d12
CD5+ d7
CD5+ d12
CD5+ d18
CD1a+ d12
CD1a+ d18
DP
CD4+ SP
CD8+ SP
E
Laurenti et al. ETP
199
5
245
Early Thymic
F
chromosome segregation
nuclear division
Cell Cycle
Cell Cycle, Mitotic
regulation of mitosis
DNA conformation change
DNA repair
DNA recombination
DNA replication
regulation of cell adhesion mediated by integrin
alpha-beta T cell activation
T cell activation
regulation of T cell differentiation
positive regulation of phospholipase C activity
Thyroid hormone synthesis
memory
G
response to interferon-gamma
lysosome
Antigen processing and presentation
positive regulation of T cell activation
MHC class II protein complex
Cytokine Signaling in Immune system
heterotrimeric G-protein complex
Primary immunodeficiency
T cell differentiation
regulation of lymphocyte activation
macrophage activation
chemokine production
myeloid leukocyte differentiation
myeloid leukocyte cytokine production
regulation of adaptive immune response
neutrophil activation
monocyte differentiation
cellular response to thyroid hormone stimulus
hematopoietic progenitor cell differentiation
secretory granule
Hematopoietic cell lineage
Transcriptional misregulation in cancer
T-helper 1 type immune response
dendritic cell differentiation

**Supplementary Figure S2: Transcriptional profiling of human thymic subpopulations. (A)** Flow cytometry sorting strategy for the indicated subpopulations isolated from human neonatal thymi. **(B-C)** Transcriptional profiling of CD34+ CD1a- CD7- (n=3), CD34+CD1a-CD7+ (n=2), CD34+CD1a+ (n=3), CD4+ISP (n=2), Double Positive (DP) T-receptor (TR) Low (n=2), DP TR High (n=3) populations sorted as in **(A). (B)** number of differentially expressed genes (FDR<0.05 by limma) in pairwise comparisons between indicated populations. **(C)** PCA of indicated populations (based on most variable genes across all thymic populations, see methods). **(D)** Comparison of the gene expression profiles of the thymic populations analyzed here with those of T-cells generated *in vitro* from CB CD34+ cells (5) on a 2D PCA. **(E)** Venn diagram showing overlap of genes in the Laurenti *et al*. ETP geneset (6) and the Early Thymic geneset identified here. Only 5 genes were found in both genesets: *MX1, LGMN, IRF8, CXCR3* and *OAS2*. **(F)** and **(G)** ClueGO pathway analysis of genes unique to **(F)** the Laurenti *et al* ETP signature and **(G)** the Early Thymic geneset identified here. Only genesets with FDR<0.05 are shown.

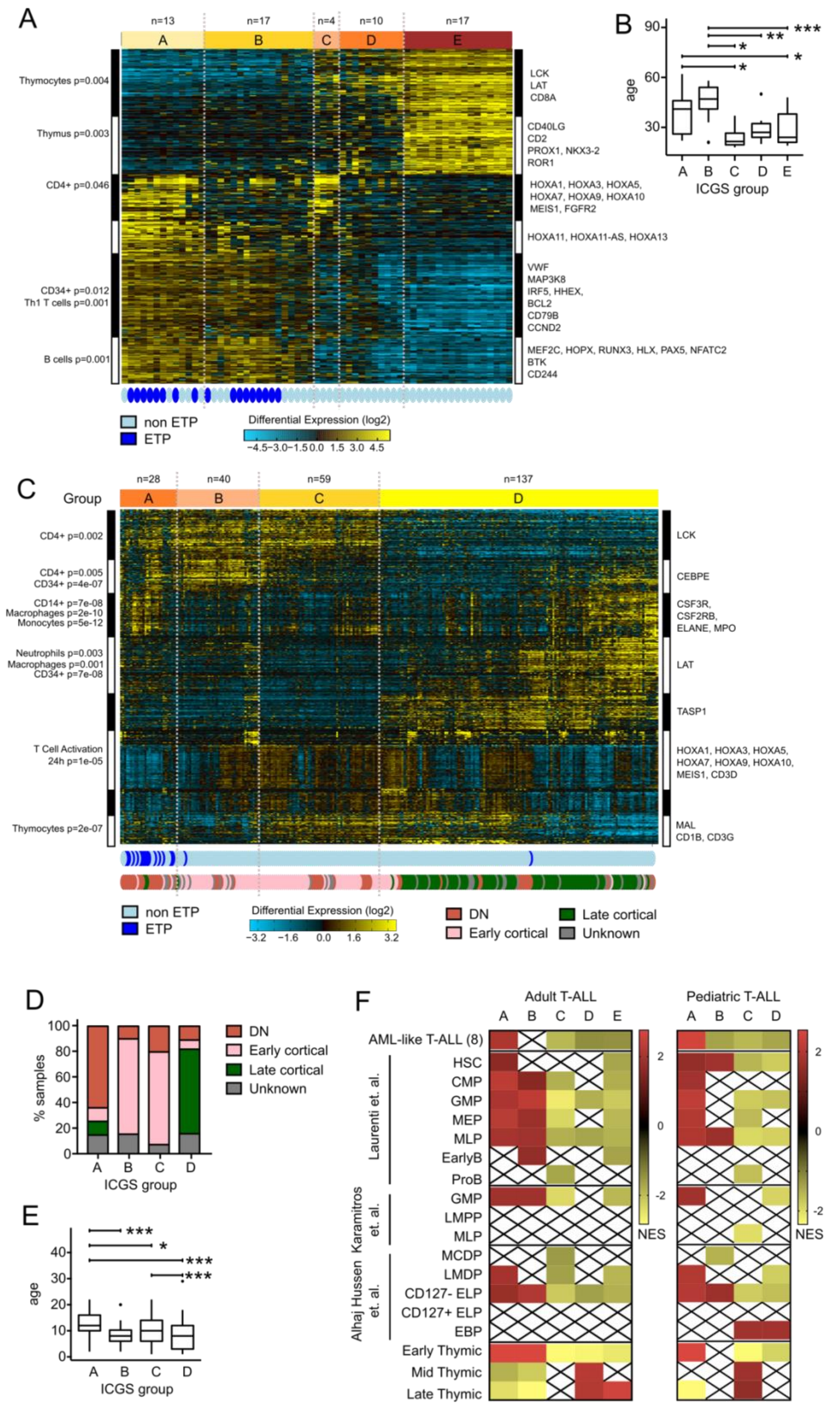

A
n=13
n=17
n=4
n=10
n=17
A
B
C
D
E
Thymocytes p=0.004
Thymus p=0.003
CD4+ p=0.046
CD34+ p=0.012
Th1 T cells p=0.001
B cells p=0.001
LCK
LAT
CD8A
CD40LG
CD2
PROX1, NKX3-2
ROR1
HOXA1, HOXA3, HOXA5,
HOXA7, HOXA9, HOXA10
MEIS1, FGFR2
HOXA11, HOXA11-AS, HOXA13
VWF
MAP3K8
IRF5, HHEX,
BCL2
CD79B
CCND2
MEF2C, HOPX, RUNX3, HLX, PAX5, NFATC2
BTK
CD244
non ETP
ETP
Differential Expression (log2)
−4.5 −3.0 −1.5 0.0 1.5 3.0 4.5
B
age
ICGS group
C
Group
n=28
n=40
n=59
n=137
CD4+ p=0.002
CD4+ p=0.005
CD34+ p=4e-07
CD14+ p=7e-08
Macrophages p=2e-10
Monocytes p=5e-12
Neutrophils p=0.003
Macrophages p=0.001
CD34+ p=7e-08
T Cell Activation
24h p=1e-05
Thymocytes p=2e-07
LCK
CEBPE
CSF3R,
CSF2RB,
ELANE, MPO
LAT
TASP1
HOXA1, HOXA3, HOXA5,
HOXA7, HOXA9, HOXA10,
MEIS1, CD3D
MAL
CD1B, CD3G
Differential Expression (log2)
−3.2 −1.6 0.0 1.6 3.2
DN
Early cortical
Late cortical
Unknown
D
% samples
ICGS group
E
age
ICGS group
F
Adult T-ALL
Pediatric T-ALL
AML-like T-ALL (8)
Laurenti et. al.
HSC
CMP
GMP
MEP
MLP
EarlyB
ProB
Karamitros et. al.
GMP
LMPP
MLP
Alhaj Hussen et. al.
MCDP
LMDP
CD127- ELP
CD127+ ELP
EBP
Early Thymic
Mid Thymic
Late Thymic
NES

**Supplementary Figure S3: ICGS classifies T-ALL samples based on their stage of leukemic differentiation arrest. (A-B)** ICGS analysis of adult samples in the Chen *et al.* dataset (1) (n=61 samples, pediatric samples were excluded). **(A)** Heatmap of expression of guide genes selected by ICGS. Genes are represented in rows. White and black bars on the side represent blocks of correlated genes and selected enriched gene ontology groups for these genes are shown. Columns represent individual T-ALL samples. Bottom bars indicate the phenotype of each individual sample, top bar indicates the clusters identified by ICGS. **(B)** Age distribution in indicated ICGS groups. * $p<0.05$, ** $p<0.01$ and *** $p<0.001$ by one-way ANOVA with multiple comparisons. **(C-E)** ICGS analysis of pediatric samples in the Liu *et al.* dataset (2) (n=264 samples). **(C)** Heatmap of expression of guide genes selected by ICGS. ETP phenotype and stage of differentiation arrest (as determined by mutation analysis in (2)) are indicated in the bottom bars, top bar indicates the major clusters identified by ICGS. **(D)** Distribution of stages of differentiation arrest in indicated ICGS groups. **(E)** Patient age distribution in indicated ICGS groups. * $p<0.05$ and *** $p<0.001$ by one-way ANOVA with multiple comparisons. **(F)** Enrichment of indicated normal hematopoietic progenitor transcriptional signatures by GSEA. Samples in the indicated ICGS clusters were compared to all other samples in each cohort. NES = Normalized Enrichment Score, crossed out boxes indicate genesets that are not significantly enriched (FDR > 0.05).

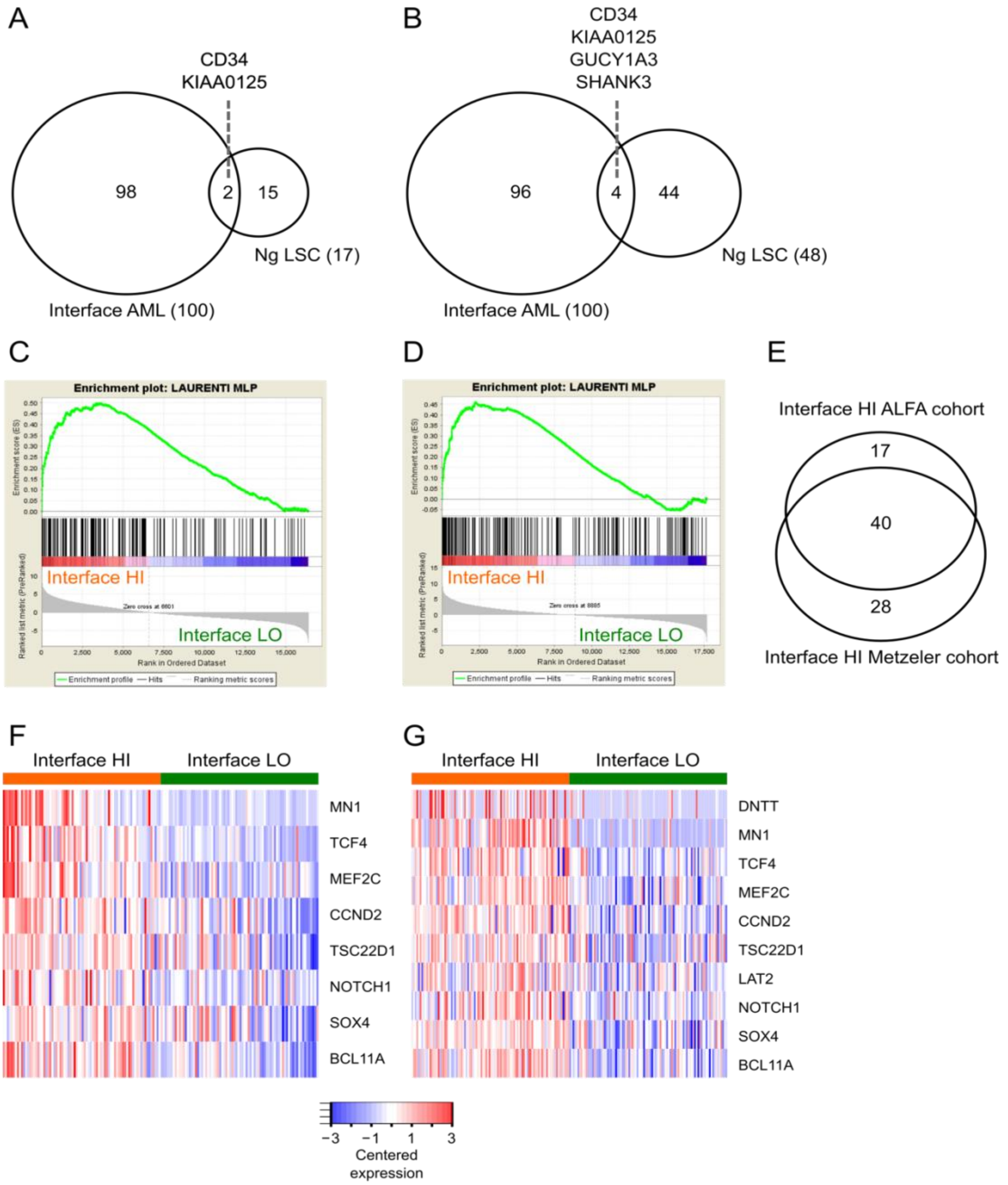


**Supplementary Figure S4: AMLs with high IAL scores are distinct from LSC-AML and are enriched for lymphoid transcriptional signatures. (A)** Overlap of IAL score with LSC17 score and **(B)** the extended 48 gene list from the same publication (7). Enrichment of MLP signatures (6) in Interface High cases by GSEA in the **(C)** Metzeler *et al* (8) and **(D)** ALFA-1701 (3) studies. These cohorts had overlap in significant differential expression of B-lymphoid genes from the MLP signature, as shown in the Venn diagram in **(E)** and in the heatmaps in **(F)** Metzeler *et al* (8) and **(G)** ALFA-1701 (3).

**Supplementary Methods:**

***Microarray experiments:*** RNA was extracted from acute leukemia and normal thymic samples using either the RNeasy Micro or Mini Kits (Qiagen), depending on cell numbers. Biotinylated double strand cDNA targets were prepared from 0.3 to 35 ng of total RNA using the NuGEN Ovation Pico WTA System V2 Kit (Cat # 3302) followed by the NuGEN Encore Biotin Module Kit (Cat # 4200) according to manufacturer recommendations. Following fragmentation, 4.5 µg of cDNAs were hybridized for 16 hours at 45$_0$C, 60 rpm on Human GeneChip® HG-U133 plus 2.0 arrays (Affymetrix). The chips were washed and stained in the GeneChip® Fluidics Station 450 (Affymetrix) using the FS450_0004 script and scanned with the GeneChip® Scanner 3000 7G (Affymetrix) at a resolution of 1.56 µm. Raw data (.CEL Intensity files) were extracted from the scanned images using the Affymetrix GeneChip® Command Console (AGCC) version 4.1.2. CEL files were further processed with Affymetrix Expression Console software version 1.4.1 to calculate probeset signal intensities, using Robust Multi-array Average (RMA) algorithms with default settings.

***Next-generation sequencing:*** Nextera XT (Illumina) DNA Libraries were prepared according to the manufacturer's instructions and sequenced using the Illumina MiSeq sequencing system. The custom NGS panel (Supplementary Table S4) was originally inspired by the repertoire of genes found to be preferentially altered in pediatric ETP-ALL (9), and we have previously reported analyses of other T-ALL cohorts using this panel (10, 11). Sequencing reads were analyzed using institutional software for alignment and mutation calling (Polyweb, Institut Imagine, Paris). Variant calling required ≥50 total reads including ≥10 alternative reads and additional visual confirmation in Integrative Genomics Viewer (https://software.broadinstitute.org/software/igv/). Variants were further filtered by reference to both constitutional (dbSNP, https://www.ncbi.nlm.nih.gov/snp/, ExAC, http://exac.broadinstitute.org/, The 1000 Genomes Browser, https://www.internationalgenome.org/1000-genomes-browsers/) and somatic (COSMIC https://cancer.sanger.ac.uk/cosmic) databases, and by prediction of mutational effect using the Polyphen (http://genetics.bwh.harvard.edu/pph2/), SIFT (https://sift.bii.a-star.edu.sg/) and Cadd (https://cadd.gs.washington.edu/score) tools.

***Data comparison with published datasets:*** Expression data from the LT-HSC, MLP, CMP, GMP, MEP, earlyB, proB umbilical cord blood (CB) populations from Laurenti *et al*(6) (GSE42414) were used to determine a list of highly variable genes across all umbilical cord blood populations (CB-HVGs, 7271 genes, defined as genes differentially expressed between any 2 populations). Microarray data from these genes was extracted from the AML/T-ALL dataset and was combined with the selected samples from GSE42414, batch corrected (using the *ComBat* function from the *sva* package) and normalised (*normalize.quantiles* function from the preprocessCore v1.34.0 package). PCA analysis was performed as above on these CB-HVGs for normal CB populations and T-ALL samples. Expression data from samples in Cante-Barret *et al* (5) (GSE79379) was combined with data from thymic populations, AMLs and T-ALLs profiled here and batch corrected (using the *ComBat* function from the *sva* package). Combined PCA analysis was performed as described in the main methods section.

***Isolation of thymic subpopulations:*** Informed consent was given for provision of normal human thymi removed during neonatal cardiac surgery at Hôpital Necker-Enfants Malades. Mononuclear cell suspensions were obtained by dissection and irrigation of thymic tissue, followed by Ficoll gradient centrifugation. Subpopulations were isolated by fluorescence activated cell sorting (FACS Aria, Becton Dickinson) using the strategy shown in Supplementary Figure S3. For thymic subpopulations CD34+CD1a-CD7-, CD34+CD1a-CD7+, CD34+CD1a+ and CD4+ ISP, FACS sorting was preceded by CD3+ and CD7+ cell depletion using magnetic activated cell sorting (MACS, Miltenyi Biotec).

***Calculation of IAL score****:* An AML interface leukemia probeset was defined as the top 100 probes (ranked by t statistic) differentially expressed between AML samples of ICGS cluster 2 and 3 (interface AMLs) and those in ICGS clusters 4 and 5. The interface score for each sample in each of the 2 independent cohorts (3, 8) was calculated as the Sum of mean-centered log2 values for each of the genes in the AML interface leukemia probeset (71 genes in (8), 82 in (3)). Patient samples in each cohort were split into interface HI and interface LO based on whether their individual interface score was above or below a threshold, set as the median of the interface score for that cohort (median interface score: -5.12 in (8), -10.17 in (3). The robustness of the approach was tested by varying this threshold: similar prognostic results were obtained for any threshold between -15 and 6 for (8), and -80 and 92 in (3).

***Outcome analyses:*** Post-hoc analyses of Overall (OS) and Event-free survival (EFS) of patients with AML were performed on publicly available data (8) and on the ALFA-1701 cohort that we have previously reported (3). Statistical analyses and survival curves were calculated in R with the *survival* package functions *survfit*, *survdiff* and *coxph* (Cox model). For clinicobiological comparisons of IAL High and Low cases in the ALFA-1701 cohort, ELN risk was defined according to the ELN-2010 classification (12), and secondary AML-type genes were defined as previously reported (4). Covariates selected for multivariate analyses (Table 1) were selected based on the results of univariate analyses (Supplementary Table S10), with additional retention of GO (gemtuzumab ozogamicin) treatment arm.

***Data availability:*** Gene expression data reported here is available at GEO as superseries GSE131207 (normal thymic populations dataset: GSE131180; AMLs/T-ALLs dataset: GSE131184).
